# Eleven biosynthetic genes explain the majority of natural variation for carotenoid levels in maize grain

**DOI:** 10.1101/2020.07.15.203448

**Authors:** Christine H. Diepenbrock, Daniel C. Ilut, Maria Magallanes-Lundback, Catherine B. Kandianis, Alexander E. Lipka, Peter J. Bradbury, James B. Holland, John P. Hamilton, Edmund Wooldridge, Brieanne Vaillancourt, Elsa Góngora-Castillo, Jason G. Wallace, Jason Cepela, Maria Mateos-Hernandez, Brenda F. Owens, Tyler Tiede, Edward S. Buckler, Torbert Rocheford, C. Robin Buell, Michael A. Gore, Dean DellaPenna

**Author notes:** Present addresses: Nacre Innovations, Houston, TX 77002 (C.B.K.); University of Illinois at Urbana-Champaign, Department of Crop Sciences, Urbana, IL 61801 (A.E.L.); University of Michigan, Ann Arbor, MI 48109 (E. W.); CONACYT - Unidad de Biotecnologia, Centro de Investigación Científica de Yucatan, Merida, Yucatan, Mexico 97200 (E. G.-C.); University of Minnesota, Bioinformatics and Computational Biology, Minneapolis, MN 55455 (J. C.); Bayer, Stonington, IL 62567 (M. M.-H.); BASF, Dawson, GA 39842 (B.F.O.); Corteva Agriscience, St. Paul, MN 55108 (T.T.).

## Abstract

Vitamin A deficiency remains prevalent in parts of Asia, Latin America, and sub-Saharan Africa where maize is a food staple. Extensive natural variation exists for carotenoids in maize grain; to understand its genetic basis, we conducted a joint linkage and genome-wide association study in the U.S. maize nested association mapping panel. Eleven of the 44 detected quantitative trait loci (QTL) were resolved to individual genes. Six of these were correlated expression and effect QTL (ceeQTL), showing strong correlations between RNA-seq expression abundances and QTL allelic effect estimates across six stages of grain development. These six ceeQTL also had the largest percent phenotypic variance explained, and in major part comprised the three to five loci capturing the bulk of genetic variation for each trait. Most of these ceeQTL had strongly correlated QTL allelic effect estimates across multiple traits. These findings provide the most comprehensive genome-level understanding of the genetic and molecular control of carotenoids in any plant system, and a roadmap to accelerate breeding for provitamin A and other priority carotenoid traits in maize grain that should be readily extendable to other cereals.

## INTRODUCTION

Carotenoids are lipid-soluble isoprenoids (typically C_40_) synthesized by plants, algae and also some fungi, bacteria and yeast (reviewed in Li *et al.* 2016). Most carotenoids have yellow, orange, or red colors that are a function of the length of their conjugated double bond system and functional groups (Khoo *et al.* 2011). Carotenoids containing oxygen functional groups are termed xanthophylls, and those without such groups are termed carotenes. In plants, carotenoids are biosynthesized and localized in plastids, where they play numerous roles in photosystem structure and light harvesting and in photoprotection through their scavenging of singlet oxygen and dissipation of excess excitation energy via the xanthophyll cycle (Jahns and Holzwarth 2012). Additionally, 9-*cis* isomers of violaxanthin and neoxanthin are precursors for biosynthesis of abscisic acid (ABA), a plant hormone with critical roles in embryo dormancy and abiotic stress responses (Kermode 2005; Tuteja 2007), while 9-*cis* β-carotene is the substrate for synthesis of strigolactones, recently discovered plant hormones involved in branching and in attracting beneficial arbuscular mycorrhizae (reviewed in Al-Babili and Bouwmeester 2015, Jia et al. 2018).

Provitamin A carotenoids are an essential micronutrient in human and animal diets, as they are converted to vitamin A (retinol) in the body via oxidative cleavage (reviewed in Eroglu and Harrison 2013). The most abundant provitamin A carotenoids in the human diet are β-carotene, which yields two molecules of retinol, and β-cryptoxanthin and α-carotene, which yield one (Stahl and Sies 2005; Combs and Mcclung 2017). Clinical vitamin A deficiency affects an estimated 127.2 million preschool children and 7.2 million pregnant women in countries determined to be at risk (West 2002). Symptoms can include xerophthalmia (‘dry eye’), which often progresses to night blindness, as well as increased morbidity and mortality from infections (reviewed in West and Darnton-hill 2008). It is estimated that vitamin A deficiency is responsible for the deaths of approximately 650,000 preschool children per year (Rice *et al.* 2004). Two non-provitamin A xanthophylls, lutein and zeaxanthin, also play important roles as macular pigments in the human fovea (Bernstein and Arunkumar 2020; Krinsky *et al.* 2003; Beatty *et al.* 1999). Elevated dietary intake of these xanthophylls have been associated with decreased risk of age-related macular degeneration (AMD; Bernstein and Arunkumar 2020; Abdel-Aal el *et al.* 2013), which affected an estimated 170 million adults in 2014 which is projected to increase as the global population ages (Wong *et al.* 2014).

Maize is a primary food staple in much of Latin America, sub-Saharan Africa, and Asia, where vitamin A deficiency remains highly prevalent (West 2002). There is extensive natural variation in levels of maize grain carotenoids, which are most highly concentrated in the vitreous (hard) portion of the endosperm (Weber 1987; Blessin *et al.* 1963; Harjes *et al.* 2008). However, as a dietary staple the average provitamin A carotenoid levels of diverse yellow maize lines provide less than 20% of the target level established based on recommended dietary allowances (Harjes *et al.* 2008; Bouis and Welch 2010; Owens *et al.* 2014). Genetic improvement of maize grain carotenoid (provitamin A) levels through breeding, an example of biofortification, has been proposed as a cost-effective approach for ameliorating vitamin A deficiency in populations who are at risk thereof (Graham *et al.* 2001; Welch and Graham 2004; Bouis and Welch 2010; Diepenbrock and Gore 2015).

Carotenoids are derived from the five-carbon central intermediate isopentenyl pyrophosphate (IPP) produced by the plastid-localized methyl-D-erythritol-4-phosphate (MEP) pathway (Figure 1). The committed step toward carotenoid biosynthesis is the head-to-head condensation of two 20-carbon geranylgeranyl diphosphate (GGDP) molecules by the enzyme phytoene synthase to form phytoene. Phytoene is sequentially desaturated and isomerized to form lycopene, which is then cyclized with two β-rings to form β-carotene, or with one β-ring and one ε-ring to form α-carotene. α- and β-carotenes are hydroxylated twice to form lutein and zeaxanthin, respectively, and zeaxanthin further modified to yield violaxanthin and neoxanthin.

**Figure 1.**
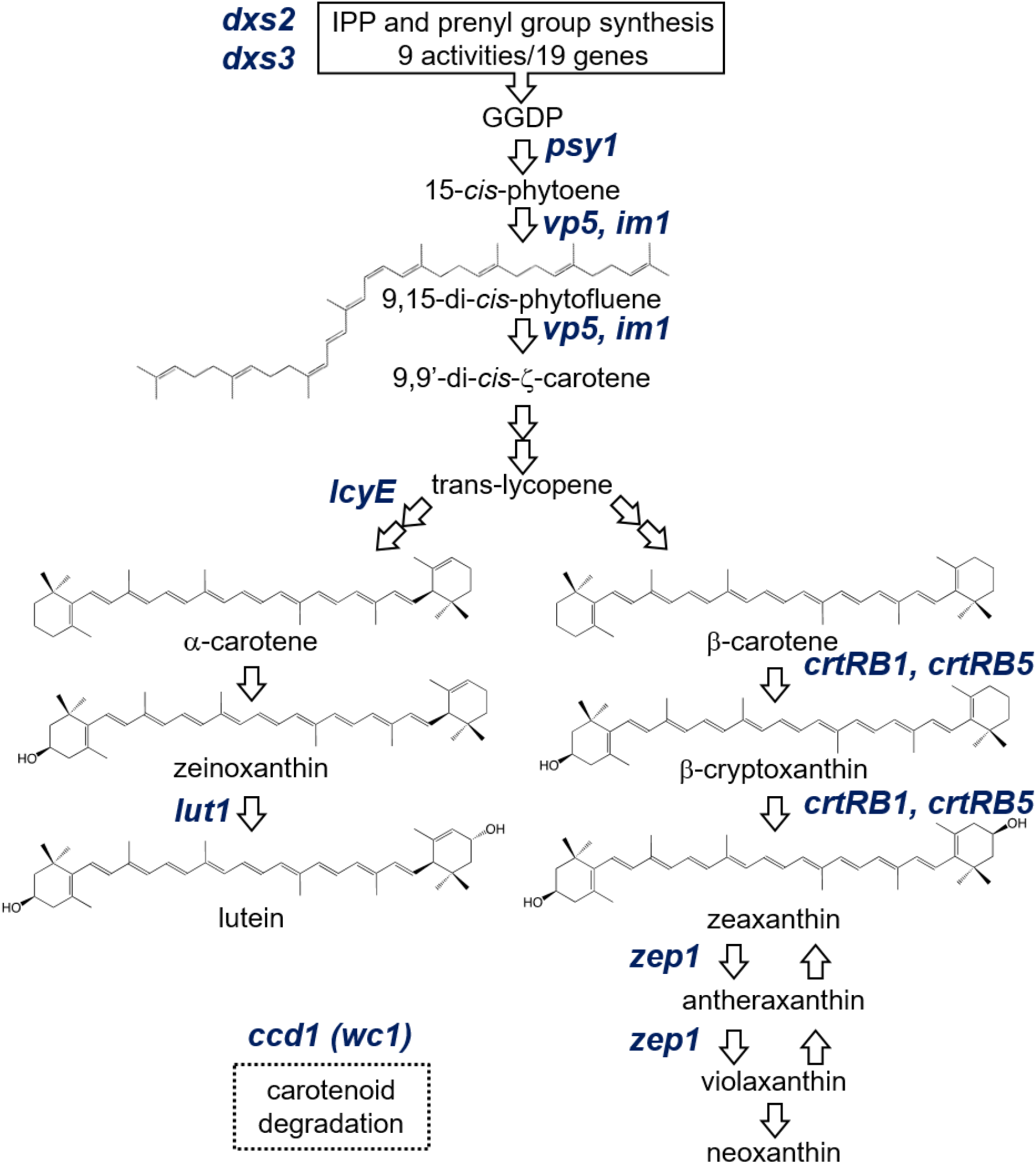
Carotenoid biosynthetic pathway in maize; grain carotenoids are primarily produced in endosperm. Precursor pathways are summarized in black boxes. The *a priori* genes identified in this study are denoted in blue italics, and are placed at the pathway step(s) executed by the enzyme that they encode. Compound abbreviation: GGDP, geranylgeranyl diphosphate. Gene abbreviations: 1-deoxy-D-xylulose-5-phosphate synthase (*dxs2* and *dxs3*); phytoene synthase (*psy1)*; phytoene desaturase (*vp5*); plastid terminal oxidase (*im1*); lycopene ε-cyclase *(lcyE);* ε-ring hydroxylase (*lut1*); β-carotene hydroxylase (*crtRB1*); zeaxanthin epoxidase (*zep1*); carotenoid cleavage dioxygenase (*ccd1*), *whitecap1* (*wc1*; a locus containing a varying number of copies of *ccd1*).

The MEP and carotenoid biosynthetic pathways are well-characterized in Arabidopsis, and the encoded genes are highly conserved across species, enabling the straightforward identification of homologs in other species. In maize, the MEP and carotenoid pathways are encoded by 58 genes, which can be considered *a priori* candidates for influencing natural variation for maize grain carotenoid levels (Supplemental Data Set 1). Four of these *a priori* candidate genes—*lycopene epsilon cyclase* (*lcyE*), *β-carotene hydroxylase 1* (*crtRB1*), *zeaxanthin epoxidase 1* (*zep1*) and *ε-ring hydroxylase* (*lut1*)—have been shown to have genome-wide associations with various carotenoid traits in maize grain (Harjes *et al.* 2008; Yan *et al.* 2010; Owens *et al.* 2014; Suwarno *et al.* 2015; Azmach *et al.* 2018; Baseggio et al. 2020). *zep1* encodes zeaxanthin epoxidase, which converts zeaxanthin to violaxanthin via antheraxanthin, and is associated with levels of zeaxanthin and total β-xanthophylls, with large explanation of phenotypic variance for these traits (Owens *et al.* 2014; Suwarno *et al.* 2015). *lut1* encodes a plastid-localized cytochrome P450 that hydroxylates the ε-ring of α-carotene to form zeinoxanthin, and is associated with zeinoxanthin levels and ratios of α-branch compounds (Owens *et al.* 2014). Lycopene epsilon cyclase (*lcyE*) catalyzes *ε*-ring cyclization of lycopene and is a key branch point enzyme: alleles with low expression in grain result in higher flux to β-carotene and β-xanthophylls (Harjes *et al.* 2008). Finally, *crtRB1* encodes a non-heme diiron hydroxylase that sequentially hydroxylates β-carotene to produce β-cryptoxanthin and zeaxanthin; weakly expressed alleles result in higher levels of β-carotene at the expense of β-xanthophylls (Yan *et al.* 2010).

Provitamin A breeding efforts in maize focusing on marker-assisted selection of favorable *lcyE* and *crtRB1* alleles have been successfully ongoing for more than a decade at two CGIAR centers, in coordination with HarvestPlus and numerous public and private partners (Saltzman *et al.* 2013; Dhliwayo *et al.* 2014; Suwarno *et al.* 2014). Maize hybrid and synthetic varieties that accumulate 40-70% of the target provitamin A level have been released, and others with higher levels are in national performance trials (Pixley *et al.* 2013, Menkir *et al.* 2017). However, introgression of favorable alleles of *lcyE* and/or *crtRB1* can have dramatically different effects depending on the genetic background (Babu *et al.* 2013; Menkir *et al.* 2017; Gebremeskel *et al.* 2018), suggesting that further investigation of these and other involved genes may facilitate and expedite the consistent achievement of target provitamin A levels. Furthermore, to simultaneously enhance and balance several other priority carotenoid traits (e.g., lutein, zeaxanthin, and total carotenoids) requires a more comprehensive understanding of the genetics underlying natural variation in maize grain carotenoid content and composition.

While substantial insight into the carotenoid pathway has been obtained from studies in Arabidopsis, its seed are green and photosynthetic whereas those of most major crops, including maize, are non-photosynthetic. Thus, maize grain is both inherently of interest as a key target crop for biofortification efforts and also potentially provides a more suitable model system for carotenoid accumulation in other major crops. In this light, we used the U.S. maize nested association mapping (NAM) panel to dissect, with high power and resolution, the quantitative trait loci (QTL) and underlying genes responsible for natural variation in grain carotenoid levels.

## RESULTS

### Genetic dissection of carotenoid accumulation in maize grain

We used the U.S. nested association mapping (NAM) panel—25 families, each comprised of 200 recombinant inbred lines (RILs) with B73 as a common parent—to dissect the genetic basis of carotenoid content and composition in maize grain. Seven grain carotenoid compounds were quantified by high-performance liquid chromatography (HPLC) with photodiode array detection. These traits and one summed trait, total carotenoids, had high estimates of line-mean heritability (0.70 to 0.94, Table 1). These traits exhibited weak negative to strong positive correlations in pairwise relationships (Supplemental Figure 1). Through a joint-linkage (JL) analysis across all 25 NAM families, we identified 117 individual-trait QTL (10 to 23 for each trait; Tables 1 and 2, Supplemental Data Sets 2 and 3) that each explained 0.7 to 40.7% of the phenotypic variance (Supplemental Data Set 4).

**Table 1.** Sample sizes, ranges, and heritabilities for carotenoid traits. Medians and ranges (in μg g^-1^ dry grain) for untransformed best linear unbiased estimators (BLUEs) of eight carotenoid grain traits evaluated in the U.S. maize nested association mapping (NAM) panel, and estimated heritability on a line-mean basis across two years. ^*a*^SD, Standard deviation of the BLUEs. ^b^Negative BLUE values are a product of the statistical analysis. Specifically, it is possible for BLUEs to equal any value from −∞ to +∞. ^*c*^SE, Standard error of the heritability estimate.

| Trait | No. Lines | BLUEs |  |  | Heritabilities |  |
| --- | --- | --- | --- | --- | --- | --- |
|  |  | Median | SD<sup>a</sup> | Range<sup>b</sup> | Estimate | SE<sup>c</sup> |
| Phytofluene | 3,521 | 0.39 | 0.54 | -0.43 - 3.02 | 0.72 | 0.02 |
| α-carotene | 3,556 | 0.91 | 0.84 | -0.70 - 4.77 | 0.70 | 0.03 |
| β-carotene | 3,538 | 0.96 | 0.97 | -0.58 - 5.49 | 0.78 | 0.03 |
| Zeinoxanthin | 3,465 | 1.44 | 1.99 | -0.95 - 11.19 | 0.89 | 0.01 |
| β-cryptoxanthin | 3,538 | 1.61 | 1.72 | -0.56 - 10.04 | 0.94 | 0.01 |
| Lutein | 3,581 | 10.68 | 6.74 | -0.72 - 39.47 | 0.94 | 0.01 |
| Zeaxanthin | 3,565 | 7.19 | 7.19 | -0.77 - 42.05 | 0.94 | 0.01 |
| Total carotenoids | 3,581 | 27.33 | 13.69 | -0.89 - 85.40 | 0.94 | 0.01 |

**Table 2.** Genetic association results for carotenoid traits. Summary of joint-linkage quantitative trait loci (JL-QTL) and genome-wide association study (GWAS) variants identified for eight carotenoid grain traits evaluated in the US maize nested association mapping (NAM) panel. ^*a*^ SD, Standard deviation. ^*b*^ GWAS variants residing within JL-QTL support intervals for each trait that exhibited a resample model inclusion probability (RMIP) of 0.05 or greater.

| Trait | Number of JL-QTL | Median size (SD<sup>a</sup>) of $\alpha = 0.01$ JL-QTL support interval (Mb) | Number of JL-QTL intervals containing *a priori* genes | Number of GWAS-associated variants in JL-QTL intervals<sup>b</sup> | Maximum RMIP |
| --- | --- | --- | --- | --- | --- |
| Phytofluene | 14 | 5.05 (27.96) | 6 | 65 | 0.67 |
| α-carotene | 11 | 2.64 (16.36) | 5 | 44 | 0.87 |
| β-carotene | 10 | 3.86 (6.72) | 8 | 39 | 0.80 |
| Zeinoxanthin | 12 | 2.42 (27.79) | 8 | 40 | 0.86 |
| β-cryptoxanthin | 23 | 3.52 (26.73) | 11 | 75 | 0.66 |
| Lutein | 14 | 3.17 (19.59) | 7 | 50 | 0.88 |
| Zeaxanthin | 18 | 4.22 (29.42) | 13 | 56 | 0.88 |
| Total carotenoids | 15 | 6.21 (20.77) | 7 | 54 | 0.73 |
| **JL-QTL TOTAL** | **117** | **3.80 (23.58)** | **65** | **423** |  |

To dissect the identified QTL at higher resolution, we performed a genome-wide association study (GWAS) using ~27 million HapMap v1 and v2 markers imputed on the ~3,600 NAM RILs (Table 2, Supplemental Data Set 5). A total of 983 marker-trait associations (101 to 142 per trait) were found to have a resample model inclusion probability (RMIP) value > 0.05 (Valdar *et al.* 2009) (Table 2, Supplemental Data Set 5). Of these, 422 (42.9%) were within a corresponding trait JL interval (Table 2), with 98 of these being attributed to 42 markers associated with two or more traits yielding a total of 366 uniquely associated markers.

Given that individual carotenoid compounds share a biosynthetic pathway (Figure 1), it was not surprising that 80% of overlapping QTL support intervals were significantly pleiotropic (Supplemental Figure 2, Supplemental Data Set 6). When the 117 individual-trait QTL intervals were merged based on physical overlap, 44 unique QTL were obtained, of which 21 impacted multiple traits (Supplemental Data Set 3). We then applied a triangulation approach (Diepenbrock *et al.* 2017) integrating JL-QTL effect estimates (Supplemental Data Set 7), GWAS marker genotypes, and RNA-sequencing (RNA-seq) expression abundances at six developing kernel stages in the NAM parents (Supplemental Data Set 8) to identify genes underlying the 44 unique QTL (Supplemental Figure 3). The rapid decay of linkage disequilibrium (LD) in proximity of GWAS-detected markers (Supplemental Figure 4), in combination with the high resolving power of the NAM panel (Wallace *et al.* 2014), supported using a search space spanning ±100 kb of those GWAS signals residing within a unique QTL for gene identification. Based on the confluence of strong triangulation correlations for a single gene within these search spaces, 11 genes were identified as underlying a unique QTL (Figure 2). All 11 genes were contained on the list of 58 *a priori* genes known from prior studies in various plants to play roles in IPP synthesis and carotenoid biosynthesis and degradation (Supplemental Data Set 1). Five of these 11 genes were in a class that we term correlated expression and effect QTL (ceeQTL), in that their expression levels were significantly associated with the JL allelic effect estimates for the QTL at multiple kernel developmental time points (Figures 2 and 3, Supplemental Figure 3). All QTL with >4% PVE were resolved down to an individual gene, with three exceptions: QTL21 for phytofluene (4.68% PVE), QTL24 for α-carotene (6.99%), and QTL34 for α-carotene (5.21% PVE) (Figure 2, Supplemental Table 1). Of the 33 JL-QTL intervals that could not be resolved to an individual gene (PVE of 0.66 to 6.99%), 22 were single-trait intervals (Supplemental Table 1).

**Figure 2.**
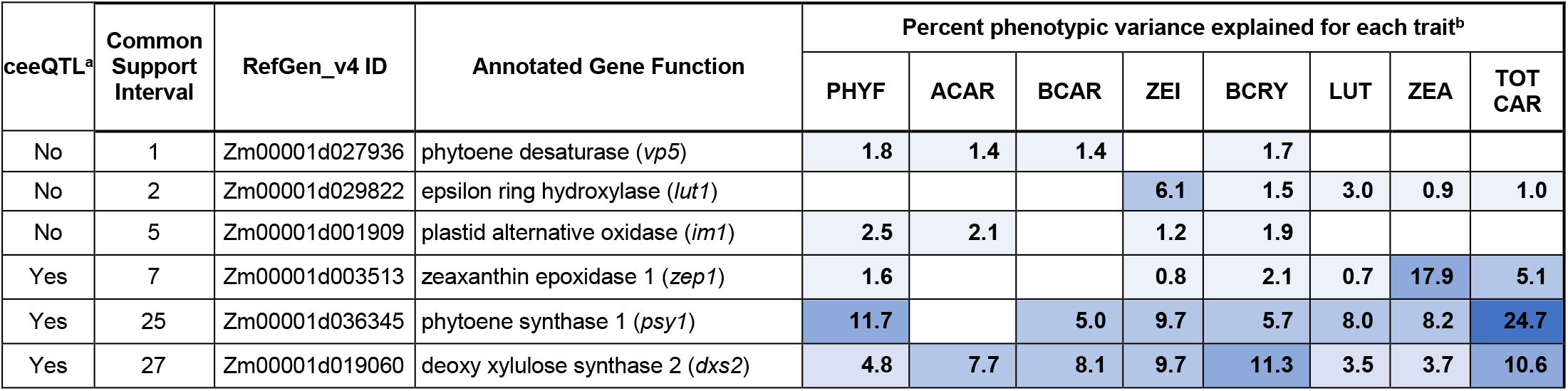
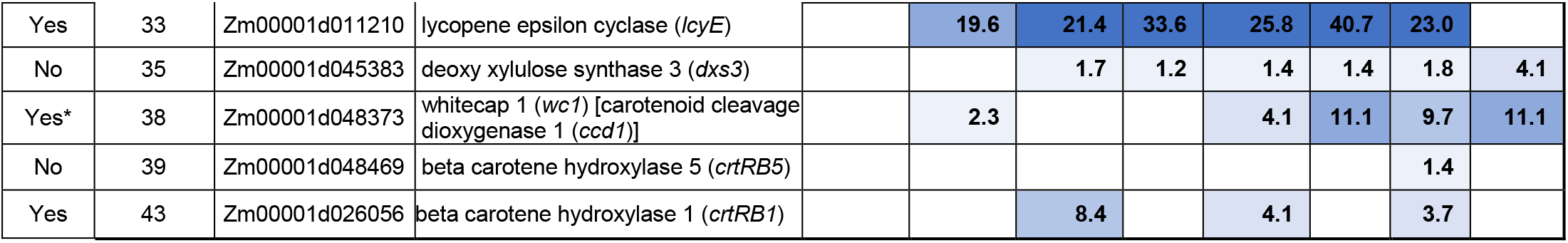
Percent phenotypic variance explained (PVE) by joint-linkage quantitative trait loci (JL-QTL). Blue shading corresponds to range of PVEs for JL-QTL, with darker blue indicating higher PVEs. ^a^ceeQTL (correlated expression and effect QTL) indicates significant correlations between expression values and JL-QTL allelic effect estimates at two or more time points for at least one trait. *The *whitecap1* locus is a macrotransposon insertion-derived, tandem-*ccd1* gene array located 1.9 Mb away from the progenitor *ccd1-r* locus (Tan et al. 2017). The high identity of mRNAs from the two loci does not allow *ccd1* mRNA from the two loci to be distinguished and therefore locus-specific FDR-corrected P-values could not be calculated. However, a strong correlation was still observed between *ccd1* expression (as well as *ccd1* copy number) and QTL38 allelic effect estimates for several traits (Figure 3, Supplemental Table 2). ^b^Trait abbreviations: PHYF, phytofluene; ACAR, α-carotene; BCAR, β-carotene; ZEI, zeinoxanthin; BCRY, β-cryptoxanthin; LUT, lutein; ZEA, zeaxanthin; TOTCAR, total carotenoids.

**Figure 3.**
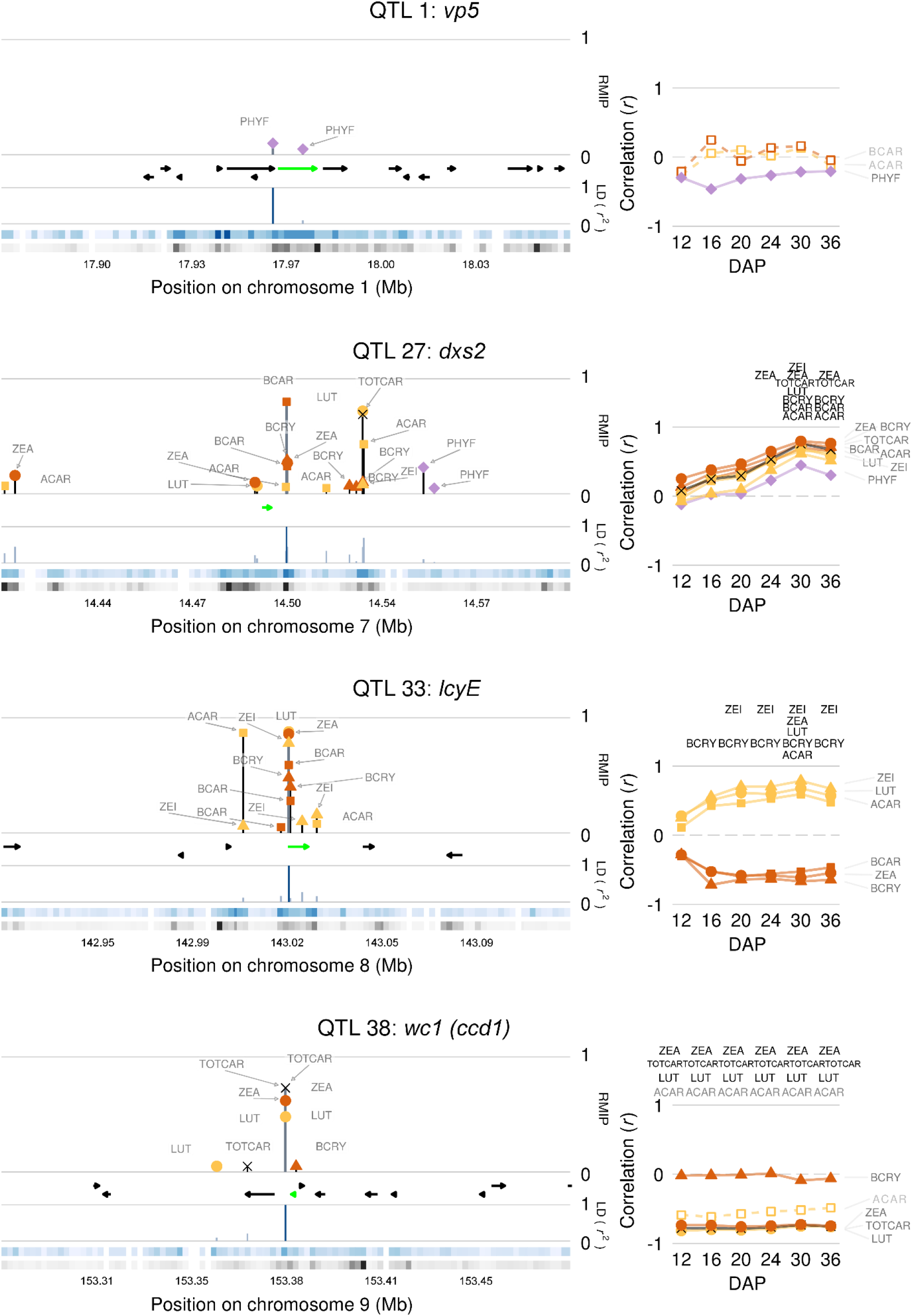
Master summaries for selected identified genes. Marker colors indicate pathway branch of the associated trait: yellow for α-branch compounds; orange for β-branch compounds; purple for phytofluene; and black for total carotenoids. Marker shapes correspond to hydroxylation state of the associated trait: squares for α- and β-carotene; triangles for β-cryptoxanthin and zeinoxanthin; circles for lutein and zeaxanthin; diamonds for phytofluene; and an X for total carotenoids. *Left panels:* Directional gene models are depicted as black arrows and the identified gene as a green arrow. Lines with trait names above gene models indicate resample model inclusion probability (RMIP) of significant GWAS hits ± 100 kb of the peak RMIP variant. Lines below gene models indicate pairwise linkage disequilibrium (LD; *r*^2^) of each GWAS variant with the peak RMIP variant (dark blue line). The blue ribbon depicts the highest LD, per 200-bp window, to the peak RMIP variant while black ribbons indicate the density of variants tested in GWAS in the 200-bp window (log_2_ scale). Darker colors correspond to higher values. *Right panels:* Correlations (*r*) between JL-QTL allelic effect estimates and expression of the identified gene across six developing kernel time points. Significant correlations are indicated by trait abbreviations above the respective time point. Traits with both JL and GWAS associations appear in black text to the right of the graph and have solid trend lines and symbols, while those with only JL associations are in gray with dashed trend lines and open symbols.

### Examination of specific variants

Two types of variant annotation analyses were employed to provide further functional insight into the 55 unique GWAS variants (i.e., 50 SNPs and five indels) involved in one or more marker-trait associations that were within ± 100 kb of the 11 identified genes: genomic evolutionary rate profiling (GERP; Davydov et al. 2010) and SNP effect analysis (SnpEff; Cingolani et al. 2012). GERP uses multi-species alignments to predict the impact of a variant as it relates to a quantitative measure of evolutionary conservation, with a positive GERP score (> 0) indicating that a site may be under evolutionary constraint and mutations with larger scores more probable to be deleterious (Davydov et al. 2010; Rodgers-Melnick et al. 2015). GERP scores from two prior studies (Kistler *et al.* 2018; Ramstein *et al.* 2019) were collectively available for 53 of the 55 unique GWAS variants. Of these, 26 variants had positive GERP scores in Kistler et al. (2018), and 12 SNPs had positive GERP scores in Ramstein et al. (2019) (Supplemental Data Set 9). SnpEff, a genomic annotation and functional effect prediction tool (Cingolani et al. 2012), predicted six of the 55 SNPs to have low or moderate effect (Supplemental Data Set 9). These results for individual genes are further detailed below.

### The role of *a priori* pathway genes

Of the eleven identified *a priori* genes, five encode enzymes that act upstream of the main carotenoid pathway branch point, where the linear carotene lycopene is cyclized to either α- or β-carotene. All five loci were associated with multiple cyclic carotenoids (Figure 2), including provitamin A compounds (β-carotene, β-cryptoxanthin, and α-carotene), and all but phytoene synthase 1 (*psy1*) represent novel associations at the genome-wide level with carotenoid variation in maize grain. Consistent with their sequential positions in the upstream portion of the pathway, all of these loci except for phytoene desaturase (*vp5*) showed positive pleiotropy (i.e., having positively correlated QTL allelic effect estimates between pairs of traits) for the downstream cyclic carotenoids with which they were associated (Supplemental File 1). Two of these five genes, *dxs2* and *dxs3*, encode 1-deoxy-D-xylulose 5-phosphate synthase (DXS), the first enzymatic reaction in the plastid-localized MEP pathway that produces IPP for biosynthesis of carotenoids and other plastidic isoprenoids. *dxs2* and *dxs3* (QTL 27 and 35, respectively) were associated with eight and six traits, with PVEs of 3.5-11.3% and 1.2-4.1%. Of the two genes, *dxs2* was found to be a strong ceeQTL in the later stages of kernel development (Figure 3). Six of 15 *dxs2* GWAS variants had positive GERP scores (Supplemental Data Set 9). Similarly, six of 12 *dxs3* GWAS variants had positive GERP scores (Supplemental Data Set 9) with one in the *dxs3* genic region (Chr9:20,245,139 bp) predicted to be a missense variant with moderate effect by SnpEff (Supplemental Data Set 9). The *dxs2* and *dxs3* homologs were the only MEP pathway genes identified in this study.

Phytoene synthase (*psy*) catalyzes synthesis of phytoene, the committed biosynthetic intermediate for all carotenoids, from two molecules of GGDP (Buckner *et al.* 1996; Li *et al.* 2008). The maize genome contains three *psy* loci, and plants having a functional *psy1* allele accumulate carotenoids in endosperm and embryo whereas those homozygous for a recessive null allele lack carotenoids in endosperm, without affecting the carotenoid content of other tissues (Buckner *et al.* 1996, Li *et al.* 2008; reviewed in Gilmore 1997; Koornneef *et al.* 2002). The *psy1* locus serves as a major genetic controller of quantitative variation for carotenoids in maize endosperm, the major site of carotenoid accumulation in grain (Zhu *et al.* 2008; Z. Fu *et al.* 2013). *psy1* (QTL 25) had 5.0-11.7% PVE for the seven carotenoids analyzed and also the largest PVE observed in this study for the sum trait of total carotenoids (24.7%). When considering only the 14 families having non-white endosperm parents, indicative of functional *psy1* alleles, this locus still showed PVEs of 1.6-10.8% for the seven measured carotenoids and 22.5% PVE for total carotenoids (Supplemental Data Set 10). *psy1* was an extremely strong ceeQTL (Supplemental Figure 3), with positive correlations between expression and effect estimates for several traits throughout kernel development, both across all 25 NAM families and across the 14 families having non-white endosperm founders (Supplemental Data Sets 7 and 11). Two of the four *psy1* GWAS variants had positive GERP scores, with one in the *psy1* genic region (Chr6:85,064,130 bp) predicted by SnpEff to be a missense variant with medium effect (HapMap_v2). This variant had the second-highest GERP score observed for GWAS variants in this study (Supplemental Data Set 9) and corresponds to a previously identified marker in the fifth exon of *psy1* termed SNP7, which was significantly associated with total carotenoid levels (Z. Fu et al. 2013). SnpEff predicted a Thr to Asn substitution, which is concordant with that identified and suggested to be causal in Z. Fu et al. (2013).

The next step in the pathway involves the sequential desaturation of phytoene to phytofluene and then ζ-carotene by phytoene desaturase (PDS, encoded by *vp5*; QTL 1), which requires plastoquinone as an electron acceptor (Mayer *et al.* 1990; Norris *et al.* 1995; Brausemann *et al.* 2017). The *vp5* locus was associated with phytofluene (a desaturation intermediate) as well as α-carotene, β-carotene, and β-cryptoxanthin, with PVEs of 1.4-1.8% (Supplemental Data Set 3). One of the two *vp5* GWAS variants (1.6 kb upstream) had a positive GERP score (Supplemental Data Set 9). Another *a priori* gene (*im1*, QTL 5) encodes a homolog of the Arabidopsis plastid terminal oxidase (PTOX, IMMUTANS) gene (Carol *et al.* 1999; Aluru *et al.* 2001), which transfers electrons from PQH_2_ to molecular oxygen to replenish PQ in the absence of an active photosynthetic electron transport chain, i.e. prior to seedling greening (reviewed in Foudree *et al.* 2012). This locus was associated with phytofluene, α-carotene, β-cryptoxanthin, and zeinoxanthin, with PVEs of 1.2-2.5%. The three *im1* GWAS variants had positive GERP scores, one of which (6.6 kb distal of the gene) had the highest GERP score observed in this study (Supplemental Data Set 9).

Four biosynthetic genes downstream of the pathway branch point were associated with carotenoid traits, and all had major effects (Figure 2). Lycopene epsilon cyclase (*lcyE*) is the committed step in α-carotene synthesis (Cunningham *et al.* 1996; Bai *et al.* 2009; Cazzonelli and Pogson 2010) and is the major genetic controller of relative flux into the α- or β-carotene pathway branches (Harjes *et al.* 2008). Accordingly, *lcyE* (QTL 33) showed negative pleiotropy (i.e., having negatively correlated QTL allelic effect estimates between pairs of traits) for compounds between the two branches and positive pleiotropy for compounds within the same branch (Supplemental File 1). This locus had the largest PVE observed in this study for each of the six α- and β-branch compounds (PVE = 19.6-40.7%) and—as also reported in Harjes *et al.* (2008)—had no significant impact on total carotenoids (Figure 2). Of six *lcyE* GWAS variants, three had positive GERP scores (Supplemental Data Set 9). Two of these GERP positive SNPs were in the genic region, of which one (Chr8:143,021,025 bp) was associated with five traits in GWAS, and predicted to be a synonymous variant with low effect by SnpEff. This SNP was 386 bp downstream of a LCYE-5′ transposable element and 347 bp upstream LCYE-SNP216, two significant variants identified in a prior study (Owens et al. 2014, Harjes et al. 2008). The other GERP positive genic variant (Chr8:143,025,685 bp) was predicted by SnpEff to cause an intron splice variant with low effect and is 100 bp downstream of a previously reported significant variant, LCYE-3′ Indel (Owens et al. 2014, Harjes et al. 2008).

Subsequent hydroxylations of the ε- and β-rings of α- and β-carotenes are performed by P450s or non-heme dioxygenases, encoded by two and six genes, respectively, in the maize genome. Of these eight genes, three were associated with carotenoid traits in maize grain: *β-carotene hydroxylase 1* (*crtRB1*) and *β-carotene hydroxylase 5 (crtRB5)*, which preferentially convert β-carotene to β-cryptoxanthin and then zeaxanthin, and CYP97C (encoded by *lut1*), a cytochrome P450-type monooxygenase that preferentially hydroxylates the ε-ring of α-carotene to yield zeinoxanthin (Tian *et al.* 2004; Quinlan *et al.* 2012). The primary impact of *lut1* (QTL 2) was on the levels of zeinoxanthin (PVE=6.1%) and lutein (PVE=2.9%, Figure 2). All three distal *lut1* GWAS variants (35.6 kb and 3.2 kb upstream and 3.6 kb downstream) had positive GERP scores (Supplemental Data Set 9). *crtRB1* (QTL 43, also known as *hyd3*) had PVEs of up to 8.4% for zeaxanthin, β-carotene, and β-cryptoxanthin, with negative pleiotropic effects between its substrate, β-carotene, and its products, β-cryptoxanthin and zeaxanthin (Supplemental File 1). Four of the six *crtRB1* GWAS variants had positive GERP scores (Supplemental Data Set 9) with one in the genic region (Chr10:137,260,105 bp) predicted by SnpEff to cause an intron splice variant with low effect. This variant is 110 bp and 1.0 kb upstream of the previously characterized crtRB1-5′TE and crtRB1-3′TE markers, respectively (Owens et al. 2014, Yan et al. 2010). *crtRB5* (QTL39, also known as *hyd5*) was identified with PVE of 1.4% for only a single trait, zeaxanthin (Figure 2) and its single GWAS variant had a positive GERP score (Supplemental Data Set 9). The subsequent β-branch enzyme, *zeaxanthin epoxidase 1* (*zep1*), had positive pleiotropic effects between β-cryptoxanthin, zeaxanthin, and total carotenoids (Supplemental File 1). *zep1* (QTL 7) primarily had a large effect on its substrate, zeaxanthin, with PVE of 17.9% (Figure 2). Concordant with this large PVE for the highly abundant compound zeaxanthin, this locus also had 5.1% PVE for the sum trait of total carotenoids (Figure 2). One of two *zep1* GWAS variants (6.3 kb upstream of the gene) had a positive GERP score (Supplemental Data Set 9).

Finally, the maize genome encodes 12 different carotenoid cleavage enzymes involved in ABA and strigolactone synthesis and carotenoid degradation. However, only one, *carotenoid cleavage dioxygenase1* (*ccd1*), was associated with carotenoid levels in this study. CCD1 has been shown to be active toward multiple cyclic and linear carotenoids *in vitro* (Vogel *et al.* 2008). A single copy of *ccd1* exists in all lines at the progenitor *ccd1-r* locus, and in a limited number of lines containing the dominant *white cap1* (*wc1*) locus a variable number of tandem *ccd1* copies (*n* = 1 to 11 in the NAM founders) are found within a Tam3L transposon inserted 1.9 Mb proximal to *ccd1-r* (Tan *et al.* 2017). The QTL identified at *wc1* had PVEs of 2.3-11.1% for four compounds, and 11.1% PVE for total carotenoids, making it the second-largest effect QTL for this trait after *psy1* (Figure 2). This locus showed positive pleiotropy for all traits detected (Supplemental File 1) and was a strong ceeQTL (Figure 3). Correlations of *ccd1* copy numbers in *wc1* alleles from Tan *et al.* (2017) with NAM JL-QTL allelic effect estimates from our study were particularly strong for the most abundant carotenoids, lutein, zeaxanthin, and total carotenoids (*r* = −0.76 to −0.85), and moderate for α-carotene (−0.56) (Supplemental Table 2). Additionally, there was a strong, positive correlation detected between *ccd1* copy number and *ccd1* expression level (log_2_-transformed FPKM) (*r* = 0.80 to 0.84) across all six developing kernel stages.

### Epistatic interactions play a minor role in determining most carotenoid traits

We tested all pairs of JL-QTL peak markers for epistasis in a joint analysis of all 25 NAM families, including 11 families that each segregated for a recessive allele of *psy1* that conditions extremely low levels of carotenoids in the endosperm. In total, 14 significant epistatic interactions were found across the 25 families, but with relatively small PVEs. For instance, *psy1* has main-effect PVEs ranging from 5.0-24.7% for seven of the eight traits (Figure 3), but was an epistatic partner in only three interactions, all with PVEs <1% (Supplemental Figure 5, Supplemental Data Set 12). Only three of the 14 interaction terms had PVEs greater than 1%: *zep1* with *wc1* for zeaxanthin (1.09%), *vp5* with QTL24 for α-carotene (6.06%), and *dxs2* with QTL34 for α-carotene (1.12%).

## DISCUSSION

This study provides the most extensive dissection to date of the quantitative genetic basis of maize grain carotenoid levels. With 11 of 44 carotenoid QTL resolved to their underlying genes, the key precursor and core biosynthetic pathway genes underpinning these traits are now largely elucidated (Figure 2, Supplemental Table 1). With the exception of phytofluene, these 11 genes explain 70.3 to 90.9% of all PVE attributed to QTL for each trait (Supplemental Figure 6). Notably, eight of the 11 genes have large effects (PVE ≥4% for one or more trait), are highly pleiotropic (Supplemental File 1) and six of the eight were also ceeQTL (Figure 2). Taken together, these findings indicate pleiotropy within the carotenoid pathway may be predominantly regulated (directly or indirectly) at the level of gene expression. A previous examination of kernel gene expression in a 508-line maize association panel supports this conclusion: J. Fu et al. (2013) identified expression QTL (marker-gene expression associations) for four of our 11 identified genes (*lcyE, crtRB1, ccd1* and *vp5*), two of which (*lcyE* and *crtRB1*) are represented in the six ceeQTL that we identified. J. Fu et al. (2013) also reported significant correlations between *lcyE* and *crtRB1* expression and grain provitamin A concentrations, providing additional support for their designation as ceeQTL.

Importantly, in addition to confirming previously reported gene/trait associations, the current study has identified new and major breeding targets in the IPP pathway and early steps of the carotenoid pathway. Four of the five genes we identified that reside upstream of lycopene cyclization (*vp5, im1, dxs2*, and *dxs3*) had not previously been associated with natural variation in maize grain carotenoids at the genome-wide level (Harjes *et al.* 2008; Yan *et al.* 2010; Owens *et al.* 2014; Suwarno *et al.* 2015; Azmach *et al.* 2018; Baseggio et al. 2020). Most notably, *dxs2* had the second-largest PVEs for four traits in this study including two provitamin A carotenoids, β-cryptoxanthin and α-carotene, at 11.3% and 7.7%, respectively*. dxs2* and *dxs3* were also associated with natural variation for vitamin E related traits (Diepenbrock *et al.* 2017), which also utilize IPP in their synthesis. Like carotenoid PVEs, *dxs2* vitamin E PVEs were greater than *dxs3* and it was also a ceeQTL (FDR < 0.05), while *dxs3* was not (Figures 2 and 3, Supplemental Figure 3). Engineering and overexpression studies in Arabidopsis and *E. coli* have shown DXS is a limiting activity in the MEP pathway (Harker and Bramley 1999; Estevez *et al.* 2001) and maize *dxs2* appears to be the major genetic control point for IPP synthesis for both carotenoids and tocotrienols, and likely other plastidic isoprenoids in maize grain.

Phytoene synthase is the committed step in carotenoid synthesis and was previously associated with variation for five carotenoids in maize kernels (Z. Fu *et al.* 2013). We confirmed that *psy1* has large PVEs for these five carotenoid traits and also two others (Figure 2). Considerable haplotype-level variation is still present at the *psy1* locus in both temperate and tropical maize (Swarts *et al.* 2017), and an allelic series was seen in the present study (Supplemental Data Set 7). This suggests that explicit attention to selecting or fixing favorable haplotypes of both *psy1* and *dxs2*, which exhibited large PVEs for many of the same traits, should increase overall flux of IPP into the carotenoid pathway and further enhance gains obtained from selection on other downstream genes (e.g., *lcyE* and *crtRB1*) for provitamin A biofortification of maize.

Genes encoding phytoene desaturase (PDS, *vp5*) and the plastid alternative oxidase (PTOX, *im1*), both of which are necessary for the sequential desaturation of phytoene to ζ-carotene via phytofluene, were identified in this study. PDS introduces double bonds into its carotenoid substrates and transfers the electrons to plastoquinone (PQ), an essential co-factor for carotenoid biosynthesis, reducing it to PQH_2_ (Norris *et al.* 1995). In photosynthetic tissues PQH_2_ is efficiently re-oxidized by the photosynthetic electron transport chain (Rosso *et al.* 2006; Shahbazi *et al.* 2007; Rosso *et al.* 2009, reviewed in Foudree *et al.* 2012). However, in non-photosynthetic tissues that have an underdeveloped photosynthetic electron transport chain—e.g., germinating seedlings and developing maize grain—PTOX transfers electrons from PQH_2_ directly to molecular oxygen to regenerate PQ for additional desaturation cycles (Beyer *et al.* 1989; Mayer *et al.* 1990; Carol *et al.* 1999; Wu *et al.* 1999, reviewed in Rodermel 2002; Foudree *et al.* 2012). PTOX loss-of-function mutants in Arabidopsis negatively affect carotenoid synthesis in developing seedlings, resulting in albino sectors of vegetative tissue that hyper-accumulate the PDS substrate phytoene (Wetzel *et al.* 1994; Carol *et al.* 1999; Wu *et al.* 1999; Rosso *et al.* 2009). Maize *vp5* and *im1* were associated with the PDS desaturation intermediate phytofluene (Figure 2) but also with three downstream provitamin A active carotenoids, suggesting *vp5* and *im1* are novel targets that should be considered for breeding or metabolic engineering efforts to enhance kernel provitamin A content.

While biosynthesis is an important control point for carotenoid levels, degradation and the susceptibility to degradation must also be considered. For example, carotenoid cleavage dioxygenase 4 (CCD4) in Arabidopsis is a large-effect contributor to natural variation in seed carotenoids (Gonzalez-Jorge *et al.* 2013), primarily due to its preferential cleavage of β-carotene and epoxy-xanthophylls, which are elevated 3-7-fold in *ccd4* null mutants. Zeaxanthin epoxidase (ZEP) has an even greater impact on total carotenoids, primarily because ZEP-mediated epoxidation increases susceptibility to degradation by CCD4. A *zep* null mutant increased zeaxanthin by 40-fold, lutein, the most abundant carotenoid in wild-type Arabidopsis seed, by 2.2-fold, and total carotenoids by 5.7-fold (Gonzalez-Jorge *et al.* 2016). In maize, *zep1* and *wc1* (encoding CCD1) had analogous impacts on natural variation of zeaxanthin, lutein and total carotenoids, and both were ceeQTL in developing kernels. Unlike CCD4, maize CCD1 is cytosolic (Tan *et al.* 2003) and only has access to the outer plastid envelope but is thought to have increased access to plastid-localized carotenoids as plastid membranes lose integrity during kernel desiccation (Tan *et al.* 2017).

A major finding of this work is that though 10 to 23 QTL were detected per trait, a large percentage of variance could be explained by only two to five major-effect QTL that contributed 4 to 41% PVE depending on the trait (Figure 2). These major-effect QTL are high-impact targets for genomics-enabled biofortification and/or metabolic engineering strategies. For example, just four genes—*lcyE, crtRB1, dxs2*, and *psy1*—explained the majority of variation for the three provitamin A carotenoids. These four genes explained 75% of β-carotene variation attributed to QTL; three of the four (*lcyE, dxs2*, and *psy1*) explained 52% for β-cryptoxanthin; and just two (*lcyE* and *dxs2*) explained 64% for α-carotene (Figure 2, Supplemental Data Set 4). These findings suggest that simultaneous increases in both β-carotene and β-cryptoxanthin, and potentially α-carotene, may be achievable by incorporating appropriate alleles of this set of four major-effect genes in selection decisions.

Currently, only *crtRB1* and *lcyE* alleles are used in marker-assisted selection efforts (Prasanna *et al.* 2020). While the combination of two favorable haplotypes of these genes was previously found to decrease total carotenoids (Babu *et al.* 2013,), neither *crtRB1* or *lcyE* (separately or in interaction) were associated with total carotenoid levels in the present study (Figure 3). Rather, total carotenoids were positively impacted by *dxs2* and *psy1* alleles and negatively conditioned by specific *zep1* and *ccd1* alleles (Figure 3). Thus, the previously reported decreases in total carotenoids observed when combining favorable *lcyE* and *crtRB1* alleles in the same background can likely be circumvented by conditioning on the presence of appropriate *psy1*, *ccd1*, and *zep1* alleles. Enabling tandem selection of *lcyE* and *crtRB1* in this manner should afford considerable gains in both total and provitamin A carotenoids, given the large PVEs and allelic effect estimates of *lcyE* observed in this study in families with both temperate and tropical founders (Figure 3, Supplemental Data Set 7). In addition to provitamin A carotenoids, lutein and zeaxanthin are the most abundant carotenoids in maize grain and are themselves of direct interest in breeding for human health given their role in eye health. Three genes—*lcyE*, *psy1*, and *wc1* (*ccd1*)—explained 77.3% of lutein variation attributed to QTL, and those three along with *zep1* explained 74.1% for zeaxanthin. The data in this study indicates simultaneous improvement and balancing of provitamin A and non-provitamin A carotenoids for human should be feasible.

The high power and resolution of the U.S. maize NAM population has allowed dissection of the majority of the phenotypic variation of maize grain carotenoids, in some cases approaching the upper limits of the high heritabilities for these traits (Table 1, Supplemental Data Set 4). The extensive information provided herein—within and across genes, and within and across traits—should focus and accelerate genomics-enabled breeding and/or metabolic engineering efforts to simultaneously achieve provitamin A targets (Bouis and Welch 2010) and improve the levels of other health-beneficial carotenoids in human populations consuming maize as a staple. The genes identified in this study are also logical candidates to be assessed as potential controllers of carotenoid variation in seeds of other abundantly consumed monocot crops.

## METHODS

### Field Environments and Plant Materials for Genetic Mapping

The design of the maize (*Zea mays*) nested association mapping (NAM) population has been previously described (Yu *et al.* 2008; Buckler *et al.* 2009; McMullen *et al.* 2009). The experimental field design in 2009 and 2010 for this study—which included the NAM panel, the intermated B73 x Mo17 (IBM) family (Lee *et al.* 2002), and a 281-line inbred diversity panel (Flint-Garcia *et al.* 2005)—was conducted as described in Chandler *et al.* (2013) and Diepenbrock *et al.* (2017). In brief, 5,000 recombinant inbred lines (RILs)—25 families, with approximately 200 recombinant inbred lines (RILs) per family—were generated by crossing 25 diverse inbred lines with B73, a common parent. The 25 families of the NAM population, along with the intermated B73 × Mo17 (IBM) family (Lee *et al.* 2002) and a 281-line inbred diversity panel designed for association mapping and used herein as checks (Flint-Garcia *et al.* 2005) were evaluated in West Lafayette, IN. These evaluations were conducted at the Purdue University Agronomy Center for Research and Education in the summers of 2009 and 2010 (i.e., two environments), using standard agronomic practices. The field design for these experiments has been previously described (Chandler *et al.* 2013). In brief, a sets design was used in each environment, with a given set containing all ~200 RILs of a family or the 281-line association panel. Each set that contained a NAM family was planted in an augmented 10 × 20 incomplete block α-lattice design, with the two parental lines included as checks in each incomplete block. Each set that contained the 281-line association panel was planted in an augmented 14 × 20 incomplete block α-lattice design, with maize inbred lines B73 and Mo17 included as checks in each incomplete block. One replicate of the entire experiment of 5,481 lines from the 25 NAM families, the IBM family, and the 281-member association panel, plus repeated check lines as described above (for the family sets and 281-line panel), was grown in each of the two environments. Two fields were used in 2009 to grow this entire single replicate, and one field was used in 2010. A single experimental unit was comprised of a single inbred line planted in a one-row (3.05 m) plot, which would contain 10 plants on average. At least four plants in each plot were self-pollinated by hand. Grain from these self-pollinated ears was harvested at physiological maturity, dried to ~15% moisture content, shelled, and bulked (within the plot) to form a representative sample for carotenoid quantification.

### Carotenoid quantification

Extraction of carotenoids was conducted on ~50 ground kernels per plot, and seven carotenoid compounds—α-carotene, β-carotene, β-cryptoxanthin, lutein, phytofluene, zeaxanthin and zeinoxanthin—as well as the sum trait of total carotenoids were quantified via high-performance liquid chromatography (HPLC) as previously described (Owens *et al.* 2014), with units of μg g^−1^ seed. Carotenoids were assessed based on HPLC data passing internal quality control measures that were collected on 9,411 grain samples from 4,871 NAM and 198 IBM RILs, as well as the 850 repeated parental check lines. HPLC-generated measurements of carotenoid compounds and their respective isomers were combined for zeinoxanthin and β-cryptoxanthin, to obtain an overall value for the level of the compound. For technical replicates of the same sample, the mean value of the replicate measurements was recorded for each of the seven carotenoids; i.e., α-carotene, β-carotene, β-cryptoxanthin, lutein, phytofluene, zeaxanthin and zeinoxanthin. Initial total carotenoid values were calculated as the sum of the quantified levels for these seven compounds.

### Phenotypic data processing

The HPLC data set was further cleaned to standardize sample genotype names, as associated with experimental field location, and any samples that lacked proper field data were removed. For each NAM family dataset, we retained only samples with genotype assignments belonging to that family or the family’s parental genotypes (which were used as checks). Samples from each NAM family and their parental genotypes were categorized as ‘yellow- to orange-grain family’ (Y) or ‘white-grain family’ (W) based on the grain endosperm color phenotype of the non-B73 parent of that family. For all samples in the ‘W’ class for which it was available, the genotype at the psy1 locus, defined as the genomic region spanning chromosome 6 position 82,017,148 to 82,020,879 bp on the AGPv2 maize reference genome, was obtained from GBS SNP marker data downloaded from MaizeGDB (Portwood et al. 2019). To eliminate possible sample contamination and select the subset of samples expected to have a functional core carotenoid pathway, samples in the ‘W’ class were further classified into ‘low’ and ‘high’ carotenoid classes using Gaussian decomposition applied to the total carotenoids values for each NAM family × year combination. Gaussian decomposition was performed using the R package ‘mclust’, specifying two mixture components and one-dimensional variable variance (Scrucca *et al.* 2016, R Core Team 2018). Samples with classification uncertainty ≥ 10% were assigned to the ‘ambiguous’ class. Any samples in the ‘W’ class that switched carotenoid class assignments between years were removed from further analysis, as well as those that were assigned to the ‘low’ or ‘ambiguous’ carotenoid class in both years or that had a homozygous alternate genotype call at the *psy1* locus. Finally, samples in both ‘W’ and ‘Y’ classes were subjected to outlier analysis. Specifically, for each NAM family × year combination, a quartile analysis was performed on the total carotenoid values using the ‘boxplot.stats’ function in R and any samples with total carotenoids values at least 1.25 * IQR (interquartile range) smaller than the first quartile were marked as low total carotenoids outliers and removed from further analysis.

Following the sample-level filtering, compound measurements were set to missing for any NAM family × year × compound measurement combination that did not have at least 20 samples measured for that compound, and all samples were removed for any NAM family × year combination which did not have at least 40 remaining samples. For the remaining samples, any missing data for a given compound was assigned a value generated by random uniform sampling from the interval (0, min_measured), where min_measured is the lowest (minimum) value measured for that compound in the corresponding NAM family × year group of samples. Following this process, the total carotenoids values were recalculated to include the assigned random uniform values.

As in Diepenbrock *et al.* (2017), the IBM RILs were not included in joint-linkage (JL) analysis or the genome-wide association study (GWAS), for they exhibit a differential recombination rate (due to being intermated). However, IBM was still included in the model fitting process to generate best linear unbiased estimators along with the 25 NAM families to provide additional information regarding spatial variation within environments and potential interactions of genotype and environment.

To examine the data for phenotypic outliers, mixed linear model selection was conducted using a custom R script that calls ASRemlR version 4 (Butler *et al.* 2017). This model selection process was conducted separately for each of the eight traits. To conduct model selection, a mixed linear model was fit where the grand mean was the only fixed effect, and random effects automatically included in the base model [i.e. without being tested in model selection] were the genotypic effects (family and RIL nested within family) and baseline spatial effects (year and field nested within year). The best random structure was then identified using the Bayesian Information Criterion (BIC; Schwarz 1978). The random structures that were tested included all combinations of the following, thus representing one to five terms (or zero, if the base model were to exhibit the most favorable BIC value) to be fit as additional random effects: a laboratory effect (HPLC auto-sampler plate) and certain additional spatial effects (set nested within field within year, block nested within set within field within year, family nested within year, and RIL nested within family within year). The best residual structure was then also identified using BIC, after the best random structure had been identified and included in the mixed linear model. The residual structures that were tested, to account for potential spatial variation across rows and/or columns within each environment, were identity by year; autoregressive for range and identity for row, by field-in-year; identity for range and autoregressive for row, by field-in-year; and autoregressive (first-order, AR1 × AR1) for range and row, by field-in-year. Field-in-year represents a new factor that combines the field name and year, to enable fitting a unique error structure for each of the three fields.

From the final fitted model for each trait, phenotypic outliers with high influence were detected using the DFFITS (‘difference in fits’) criterion (Neter *et al.* 1996; Belsley *et al.* 2005) as previously described in Diepenbrock *et al.* (2017), and observations were set to NA if they exceeded a conservative DFFITS threshold previously suggested for this experimental design (Hung *et al.* 2012). Following outlier removal, the model-fitting process described above was conducted again to estimate best linear unbiased estimators (BLUEs) for the RILs, but with the genotypic effects of family and RIL within family now included as sparse fixed effects rather than random effects. Note that the model was first fit in this step with these genotypic effects as random, then updated with these effects as sparse fixed. All terms except for the grand mean were then again fitted as random effects to estimate variance components for the calculation of line-mean heritabilities. These heritability estimates were calculated only across the 25 NAM families (Hung *et al.* 2012), and the delta method was used to obtain standard errors (Holland *et al.* 2003).

The BLUEs generated for each trait were then examined to detect any remaining statistical outliers. Specifically, the Studentized deleted residuals (Kutner *et al.* 2004) were examined using PROC MIXED in SAS version 9.3 (SAS Institute 2011). These residuals were obtained as in Diepenbrock *et al.* (2017), from a parsimonious linear model that contained the grand mean and a single randomly sampled, representative SNP (PZA02014.3) from the original genetic map for the NAM panel (McMullen *et al.* 2009), as fixed effects. The BLUE value of a given RIL for a given trait was set to NA if its corresponding Studentized deleted residual had magnitude greater than the Bonferroni critical value of *t*(1 – α/2*n*; *n* – *p* – 1). The significance level (α) used in this step was 0.05; *n* was the sample size of 3,585 RILs; and *p* was the number of predictors.

The Box-Cox power transformation (Box and Cox 1964) was then performed separately on BLUEs for each trait as in Diepenbrock *et al.* (2017). Briefly, the same parsimonious model used to generate Studentized deleted residuals was also used to identify the most appropriate Box-Cox transformation (with λ tested between −2 and 2, step of 0.05) that corrected for heteroscedasticity and error terms that were not normally distributed. PROC TRANSREG within SAS version 9.3 (SAS Institute 2011) was used to find the optimal λ for each trait (Supplemental Table 3) and apply the transformation. Note that the Box-Cox power transformation requires positive input values. Each of the traits had some number of negative BLUE values (ranging from 1 to 283 RILs per trait); these are a reasonable result of the BLUE-fitting process (Burkschat 2009). The lowest possible integer needed to make all values positive for a given trait was added as a constant across that trait vector for all of the RILs before applying the transformation; this constant had a value of 1 for all traits.

### Joint linkage analysis

A 0.1 cM consensus genetic linkage map (14,772 markers) was used for joint linkage (JL) analysis, as in Diepenbrock et al. (2017). This map was generated by imputing SNP genotypes at 0.1 cM intervals in a process previously described (Ogut *et al.* 2015), using genotyping-by-sequencing (GBS) data for ~4,900 NAM RILs as anchors (Elshire *et al.* 2011; Glaubitz *et al.* 2014). JL analysis was then conducted as previously described (Diepenbrock *et al.* 2017) across the 25 families of the NAM population to map QTL for natural variation in one or more maize grain carotenoid traits. Briefly, joint stepwise regression was implemented using modified source code in TASSEL version 5.2.53 (Bradbury *et al.* 2007; modified source code provided on GitHub), with transformed BLUEs as the response variable and the family main effect forced into the model first as an explanatory variable. The effects of each of the 14,772 markers in the 0.1 cM linkage map, nested within family, were then tested for inclusion in the final model as explanatory variables. The significance threshold for model entry and exit of marker-within-family effects was based on conducting JL analysis on 1000 permutations of transformed BLUEs for each trait and selecting the entry *P*-value thresholds (from a partial F-test) that control the Type I error rate at α = 0.05. The permutation-derived entry thresholds are listed in Supplemental Table 3. Exit thresholds were set to be twice the value of these empirically derived entry thresholds, so that a marker could not enter and exit the model in the same step.

Upon examining the results of initial JL analysis conducted via the above-described procedure, peak markers in the vicinity of *psy1* for various traits, which is the only locus in this genomic interval expected to control for the presence/absence of endosperm carotenoids in white-grain families, were found to have low minor allele counts exclusively among white-grain families (with fewer than 40 individuals across all 25 families having a genotypic state score greater than zero; zero represents homozygosity for the major allele), resulting in apparent inflation of the phenotypic variance explained by these markers. The RILs that had a genotypic state score greater than zero at the *psy1*-proximal peak marker for a given trait were removed from the data set for all traits (prior to the DFFITs step and BLUE generation), comprising 54 unique RILs removed in total. The analytical pipeline was then re-conducted from the mixed linear model selection step to re-generate JL models, and for use of the results in all downstream analyses. After this additional removal step, it was confirmed that no markers in the vicinity of *psy1* exhibited significant JL signal in any white-grain families. The sample sizes per population in this final data set used in all 25-family analyses are listed in Table 1. The permutation procedure was also applied within the 14 NAM families having parents with non-white endosperm color to enable an additional, separate JL analysis within those families using appropriate thresholds, e.g. due to the reduced sample size.

Some multicollinearity between markers in the consensus map was expected, and indeed for two traits (zeaxanthin and zeinoxanthin), two pairs and one pair of markers present in the final JL model, respectively, had a Pearson’s correlation coefficient (*r*) with magnitude greater than 0.8 between their SNP genotype states. In these cases, the marker with smaller sum of squares within the JL model was removed. A re-scan procedure was then conducted in the vicinity of any of the remaining peak markers for a given trait to test whether the removal of the multicollinear marker(s) meant a shift in the association signal. Specifically, if another marker within the support interval now had a larger sum of squares than the original peak marker, that marker would replace the original peak marker in the model, and this process was repeated (including with re-calculation of the support interval) until a local maximum in the sum of squares was found. The final peak JL markers following re-scan, along with the family term, were then re-fitted to obtain final statistics from the JL analysis of each trait. Allelic effect estimates for each QTL, nested within family, were generated as in Diepenbrock *et al.* (2017) by fitting final JL models with the ‘lm’ function (R, *lme4* package), which also evaluates the significance of these QTL-within-family terms in two-sided independent t-tests. In this step, the false discovery rate was controlled at 0.05 via the Benjamini-Hochberg procedure (Benjamini and Hochberg 1995).

Support intervals (α = 0.01) were calculated for the JL-QTL in each final model as previously described (Tian *et al.* 2011). Logarithm of the odds (LOD) scores were calculated (R, ‘logLik’ base function). The phenotypic variance explained (PVE) by each joint QTL was calculated using previous methods (Li *et al.* 2011), with some modifications as described in Diepenbrock *et al.* (2017) to account for segregation distortion across the families. While transformed data were used throughout the analyses in the present study, including in all steps that required statistical inference, it was also desired to more closely examine the signs and magnitudes of QTL allelic effect estimates on the original trait scale and in directly interpretable units of nutrition. For this single purpose, the final JL model determined using transformed BLUEs was refit with untransformed BLUEs without further model selection or re-scan.

### Genome-wide association study

Chromosome-specific residuals for each trait were obtained from the final transformed JL models with the family term and any joint QTL located on the given chromosome removed. These residuals were used as the response variable in GWAS, whereas the genetic markers tested as explanatory variables consisted of the 26.9 million variants (SNPs and indels <15 bp) of the maize HapMap v. 1 and 2 projects (Gore et al. 2009, Chia et al. 2012), as previously described (Wallace *et al.* 2014), that were upliftable to RefGen_v4 coordinates. Uplifting of HapMap markers from the B73 RefGen_v2 to RefGen_v4 assembly was conducted by clipping 50 nucleotides from each side of a given marker in its v2 position (101 nucleotides of flanking sequence in total). These were then aligned to the B73 RefGen_v4 assembly using Vmatch (v2.3.0; Kurtz 2019), with the options of -d -p -complete -h1. The resulting alignments were then filtered to keep the highest scoring and unique alignment for each marker. If a marker did not have a high-confidence, unique alignment, it was omitted from the set of upliftable markers.

The upliftable markers were projected onto the NAM RILs using the dense 0.1 cM resolution linkage map, and GWAS was conducted in the NAM-GWAS plugin in TASSEL version 4.1.32 (Bradbury *et al.* 2007) as previously described (Wallace *et al.* 2014, Diepenbrock *et al.* 2017). Briefly, a forward selection regression procedure was conducted 100 times for each chromosome, with 80% of the RILs from every family sub-sampled each time. For each trait, the model entry threshold was empirically determined by conducting GWAS on 1000 permutations of chromosome-specific residuals for each trait and averaging the results across chromosomes (Wallace *et al.* 2014) to control the genome-wide Type I error rate at α = 0.05 (Supplemental Data Set 12). The significance threshold used for a marker in GWAS was its resample model inclusion probability (RMIP) value, or the proportion of the 100 final GWAS models in which that marker was included (i.e. meeting the model entry threshold). Markers having an RMIP > 0.05 were considered in downstream analyses.

### RNA sequencing

Sample collection for RNA-seq—in three biological replicates of the NAM founders at six developing kernel stages, and in root and shoot tissues—and RNA sequencing and sample quality assessment were as conducted in Diepenbrock et al. (2017). Briefly, one self-pollinated ear per plot was sampled for each developing kernel stage, frozen in liquid nitrogen in the field and held at −80 °C until shelling and the removed kernels were stored at −80 °C. Thirty kernels were then randomly sampled from across the replicates for a given parent, and bulked; for the majority of samples, 10 seeds were used per replicate. For root and shoot samples, seed were surface sterilized and germinated on wet filter paper for four to five days at room temperature under grow lamps. Germinated seedlings were then transplanted into soil in pots and placed in a long-day greenhouse for an additional 14 days at 30 to 33 °C. Plants were removed from pots, rinsed with water to remove soil, and root and shoot tissue harvested separately and flash-frozen in liquid nitrogen. Samples were stored at −80 °C until RNA extraction. Equal weights of shoots and roots were sampled from across the replicates for a given parent and bulked. Total RNA was extracted and sequenced, and raw reads processed and aligned, as previously described with no modifications (Diepenbrock et al. 2017). In the present study, RNA-seq reads from Diepenbrock *et al.* (2017) were directly downloaded from the National Center for Biotechnology Information Sequence Read Archive (BioProject PRJNA174231) and processed using the same pipeline as described in Hoopes et al. (2018). In brief, read quality was assessed using FASTQC (https://www.bioinformatics.babraham.ac.uk/projects/fastqc/) and MultiQC (Ewels *et al.* 2016) and then cleaned using Cutadapt (Martin 2011) to remove adaptors and low quality sequences, aligned to AGPv4 of B73 using TopHat2 (Kim *et al.* 2013) with the parameters -i 5 -I 60000 --library-type fr-unstranded, and expression abundances determined using Cufflinks (Trapnell *et al.* 2012) in the unstranded mode with a maximum intron length of 60 kb and the AGPv4 annotation. Expression data are available via the MSU Maize Genomics Resource (Hoopes *et al.* 2018; http://maize.plantbiology.msu.edu/index.shtml) via a JBROWSE installation and as a downloadable expression matrix.

### FPKM filtering

The gene set was filtered (as in Diepenbrock et al. 2017) such that at least one of the kernel developmental samples in at least one sampled founder line had an FPKM greater than 1.0; a total of 30,121 genes remained upon filtering with this criterion. Expression data for genes passing the specified threshold were transformed according to log_2_(FPKM + 1), where the constant of 1 was added to allow the transformation of ‘0’ values. These log_2_-transformed values are herein specified as “gene expression levels” (Supplemental Data Set 8).

### Triangulation analysis

The JL support intervals from two or more individual-trait models that were physical overlapping were merged to form common support intervals, as in Diepenbrock *et al.* (2017). Physically distinct support intervals detected for a single trait were also retained. Triangulation analyses were conducted as in Diepenbrock *et al.* (2017), based on all pairwise Pearson correlations between trait JL-QTL effect estimates; marker genotype state for each significant GWAS marker in the interval for the respective trait(s); and log_2_-transformed expression values of genes within ± 100 kb of any of these significant GWAS markers. The search space of 100 kb was selected based on LD decay (Supplemental Figure 4). For those correlations involving one of the five traits with a negative optimal lambda for the Box-Cox transformation (i.e., an inverse power transformation was applied for these traits), the sign of the correlation was reversed in graphical and tabular representations (Figure 3 and Supplemental Figure 3 for master gene summaries, Supplemental Figure 2 and Supplemental Data Set 6 for pleiotropy) to represent the true directionality of the relationship between traits.

### Epistasis

For the peak markers in the final JL model for each trait, each possible additive x additive pairwise interaction was individually tested for significance in a model containing all marker main effects as in Diepenbrock *et al.* (2017). This procedure was conducted both in all 25 families and in the 14 families having non-B73 parents with non-white endosperm. The model entry thresholds for these interaction terms was determined (separately for the 25- and 14-family data sets) by modeling 1000 null permutations of transformed trait BLUEs with only additive terms in the model, and selecting the *P*-value approximating a Type I error rate at α = 0.05. Final epistatic models were then fit with all marker main effects and any significant interactions. PVE was calculated as described above, except that pairwise genotype scores were collapsed into three classes for interaction terms as previously described (Diepenbrock *et al.* 2017). Significant interactions in the 25-family analysis were graphically depicted using the Circos software package (Krzywinski *et al.* 2009) (Figure 4).

### Pleiotropy

Pleiotropy was assessed as previously described (Buckler *et al.* 2009), by applying the JL QTL model for each trait to every other trait. Pearson correlations between the allelic effect estimates for the original trait and the trait to which its model was applied were evaluated for significance at α = 0.01 after FDR correction via the Benjamini-Hochberg method. Significant pleiotropic relationships were visualized using the *network* R package (Butts 2008; Butts 2015) (Supplemental Figure 2). Pleiotropy was also examined within each common support interval (Supplemental File 1) to validate the merging of individual-trait intervals, a step conducted in previous NAM JL analyses (Tian *et al.* 2011). In this QTL-level analysis, each peak JL marker within the interval was fit for every other trait that had a peak JL marker in the interval.

### Linkage disequilibrium analysis

The same imputed genotypic data set of 26.9 million segregating markers used in JL-GWAS was used to estimate LD. Specifically, pairwise linkage disequilibrium (LD) of each significant GWAS marker with all other markers within ± 250 kb was estimated through custom Python and R scripts as previously described (Weir 1996; Wallace *et al.* 2014; Diepenbrock *et al.* 2017). A null distribution was generated by performing the same estimation for 50,000 markers selected at random. LD was examined in both v2 and uplifted v4 coordinates; in the latter case, the small minority of flanking markers that moved outside of the ± 250 kb region upon uplifting were dropped from the v4 analysis for that given marker.

### Variant annotation

Variant annotation was conducted on the GWAS variants in this study; i.e., genetic markers that were significant for one or more traits in GWAS, and that were within ± 100 kb of an identified gene. Existing genomic evolutionary rate profiling (GERP) scores in maize were downloaded from the publicly available data sets of Ramstein et al. (2019) and Kistler et al. (2018). GERP scores [along with minor allele frequencies, from Ramstein et al. (2019)] were extracted for the GWAS variants in this study based on their AGP_v4 chromosome and position. Annotation of genetic variants and prediction of effects was conducted in SnpEff (Cingolani et al. 2012). First, the input files used in GWAS were converted to VCFs in TASSEL 5.2.65 (Bradbury *et al.* 2007). The GWAS variants in this study were then annotated using AGP_v4 coordinates in SnpEff 5.0 (build 2020-08-09) (Cingolani et al. 2012), using the following command (and Ensembl Genome release 46; ftp://ftp.ensemblgenomes.org/pub/release-46): java -Xmx8g -jar snpEff.jar Zea_mays [infile].vcf > [outfile]_Annot.vcf.

## AUTHOR CONTRIBUTIONS

C.H.D., M.A.G., and D.D.P co-wrote the manuscript; C.H.D., D.C.I., C.B.K., and A.E.L. co-led data analysis; M.M.-L. performed HPLC analyses and metabolite quantifications; B.V., E.G.-C., J.P.H., E.W., and J.C. performed transcriptome analysis; J.P.H. performed uplifting analyses; J.G.W. generated marker data sets; J.G.W. and D.C.I. coded GWAS, GWAS permutation, and figure scripts; J.C. created website/databases; D.C.I. and P.J.B co-led phenotypic data processing; J.B.H. conceptualized and coded spatial model fitting and heritability implementation; P.J.B. modified TASSEL source code and oversaw epistasis calculations; T.R. overall management of NAM population growth; M.M.-H. managed planting, pollination, harvesting, processing of NAM population; B.F.O. and T.T. generated developing kernels for transcriptome analysis; E.S.B. oversaw germplasm, genotyping, and imputation, and advised on mapping analysis; C.R.B. oversaw RNA-seq and transcriptome analysis, managed all informatics; M.A.G. oversaw data analysis, project management, design, coordination; D.D.P. overall project management and coordination, oversaw data and metabolite analyses, biological interpretation.

## ACKNOWLEDGMENTS

This research was supported by the National Science Foundation (DBI-0922493 to D.D.P., C.R.B, E.S.B., and T.R. and DBI-0820619 and IOS-1238014 to E.S.B.), by the USDA-ARS (E.S.B.), by Cornell University startup funds (M.A.G.), and by the University of California, Davis startup funds (C.H.D.). We gratefully acknowledge Dr. Arthur Gilmour for expert support in AsREML, and Dr. Guillaume Ramstein for advisement related to GERP scores.

## REFERENCES

Abdel-Aal el, S. M., H. Akhtar, K. Zaheer and R. Ali, 2013 Dietary sources of lutein and zeaxanthin carotenoids and their role in eye health. Nutrients 5: 1169–1185.

Al-Babili, S., and H. J. Bouwmeester, 2015 Strigolactones, a novel carotenoid-derived plant hormone. Annu Rev Plant Biol 66: 161–186.

Aluru, M. R., H. Bae, D. Wu and S. R. Rodermel, 2001 The Arabidopsis *immutans* mutation affects plastid differentiation and the morphogenesis of white and green sectors in variegated plants. Plant Physiol. 127: 67–77.

Azmach, G., A. Menkir, C. Spillane, and M. Gedil 2018 Genetic loci controlling carotenoid biosynthesis in diverse tropical maize lines. G3: Genes|Genomes|Genetics 8: 1049–1065.

Babu, R., N. P. Rojas, S. Gao, J. Yan and K. Pixley, 2013 Validation of the effects of molecular marker polymorphisms in LcyE and CrtRB1 on provitamin A concentrations for 26 tropical maize populations. Theor. Appl. Genet. 126: 389–399.

Bai, L., E. H. Kim, D. DellaPenna and T. P. Brutnell, 2009 Novel lycopene epsilon cyclase activities in maize revealed through perturbation of carotenoid biosynthesis. Plant J. 59: 588–599.

Baseggio, M., M. Murray, M. Magallanes-Lundback, N. Kaczmar, J. Chamness et al., 2020 Natural variation for carotenoids in fresh kernels is controlled by uncommon variants in sweet corn. Plant Genome 13(1): e20008.

Beatty, S., M. Boulton, D. Henson, H.-H. Koh and I. J. Murray, 1999 Macular pigment and age related macular degeneration. Br. J. Ophthalmol. 83: 867–877.

Belsley, D. A., E. Kuh and R. E. Welsch, 2005 Regression Diagnostics: Identifying Influential Data and Sources of Collinearity. John Wiley & Sons, Hoboken, New Jersey.

Benjamini, Y., and Y. Hochberg, 1995 Controlling the false discovery rate: a practical and powerful approach to multiple testing. J. Royal Stat. Soc. B Met. 57: 289–300.

Bernstein, P. S., and R. Arunkumar, 2020 The emerging roles of the macular pigment carotenoids throughout the lifespan and in prenatal supplementation. J. Lipid Res.: jlr.TR120000956.

Beyer, P., M. Mayer and H. Kleinig, 1989 Molecular oxygen and the state of geometric isomerism of intermediates are essential in the carotene desaturation and cyclization reactions in daffodil chromoplasts. Eur. J. Biochem. 184: 141–150.

Blessin, C. W., J. D. Brecher, and R. J. Dimler, 1963 Carotenoid of corn and sorghum: V. Distribution of xanthophylls and carotenes in hand-dissected and dry-milled fractions of yellow dent corn. Cereal Chem. 40: 582–586.

Bouis, H. E., and R. M. Welch, 2010 Biofortification--a sustainable agricultural strategy for reducing micronutrient malnutrition in the Global South. Crop Sci. 50: S-20–S-32.

Box, G. E. P., and D. R. Cox, 1964 An analysis of transformations. J. Royal Stat. Soc. B Met. 26: 211–252.

Bradbury, P. J., Z. Zhang, D. E. Kroon, T. M. Casstevens, Y. Ramdoss et al., 2007 TASSEL: software for association mapping of complex traits in diverse samples. Bioinformatics 23: 2633–2635.

Brausemann, A., S. Gemmecker, J. Koschmieder, S. Ghisla, P. Beyer et al., 2017 Structure of phytoene desaturase provides insights into herbicide binding and reaction mechanisms involved in carotene desaturation. Structure 25: 1222–1232 e1223.

Brown, P. J., N. Upadyayula, G. S. Mahone, F. Tian, P. J. Bradbury et al., 2011 Distinct genetic architectures for male and female inflorescence traits of maize. PLOS Genet. 7: e1002383.

Buckler, E. S., J. B. Holland, P. J. Bradbury, C. B. Acharya, P. J. Brown et al., 2009 The genetic architecture of maize flowering time. Science 325: 714–718.

Buckner, B., P. S. Miguel, D. Janick-Buckner, and J. L. Bennetzen, 1996 The y1 gene of maize codes for phytoene synthase. Genetics 143(1): 479–488.

Burkschat, M., 2009 Linear Estimators and Predictors Based on Generalized Order Statistics from Generalized Pareto Distributions. Commun. Stat.-Theor. M. 39: 311–326.

Butler, D. G., Cullis, B.R., A. R. Gilmour, Gogel, B.G. and Thompson, R. 2017. ASReml-R Reference Manual Version 4. VSN International Ltd, Hemel Hempstead, HP1 1ES, UK.

Butts, C., 2008 Network: a package for managing relational data in R. J. Stat. Softw. 24.

Butts, C., 2015 Network: Classes for Relational Data, in The Statnet Project (http://statnet.org).

Carol, P., D. Stevenson, C. Bisanz, J. Breitenbach, G. Sandmann et al., 1999 Mutations in the Arabidopsis gene *IMMUTANS* cause a variegated phenotype by inactivating a chloroplast terminal oxidase associated with phytoene desaturation. Plant Cell 11: 57–68.

Cazzonelli, C. I., and B. J. Pogson, 2010 Source to sink: regulation of carotenoid biosynthesis in plants. Trends Plant Sci. 15: 266–274.

Chandler, K., A. E. Lipka, B. F. Owens, H. Li, E. S. Buckler et al., 2013 Genetic analysis of visually scored orange kernel color in maize. Crop Sci. 53: 189–200.

Chen, Y., J. Li, K. Fan, Y. Du, Z. Ren et al., 2017 Mutations in the maize zeta-carotene desaturase gene lead to viviparous kernel. PLOS ONE 12: e0174270.

Chia, J. M., C. Song, P. J. Bradbury, D. Costich, N. de Leon et al., 2012 Maize HapMap2 identifies extant variation from a genome in flux. Nat. Genet. 44: 803–807.

Cingolani, P., A. Platts, L. L. Wang, M. Coon, T. Nguyen et al., 2012 A program for annotating and predicting the effects of single nucleotide polymorphisms, SnpEff: SNPs in the genome of *Drosophila melanogaster* strain *w*^1118^*; iso*-2; *iso*-3. Fly 6(2): 80–92.

Combs, G. F., and J. P. McClung, 2017 Vitamin A. The vitamins: Fundamental aspects in nutrition and health, 5th Edition, Academic Press, London: 93–138.

Cunningham, F. X., Jr., B. J. Pogson, Z. Sun, K. A. McDonald, D. DellaPenna et al., 1996 Functional analysis of the β and ε lycopene cyclase enzymes of Arabidopsis reveals a mechanism for control of cyclic carotenoid formation. Plant Cell 8: 1613–1626.

Davydov, E. V., D. L. Goode, M. Sirota, G. M. Cooper, A. Sidow, and S. Batzoglou, 2010. Identifying a high fraction of the human genome to be under selective constraint using GERP++. PLOS Comput. Biol. 6(12): e1001025.

Dhliwayo, T., N. Palacios-Rojas, J. Crossa and K. V. Pixley, 2014 Effects of S1 recurrent selection for provitamin A carotenoid content for three open-pollinated maize cultivars. Crop Sci. 54: 2449–2460.

Diepenbrock, C. H., C. B. Kandianis, A. E. Lipka, M. Magallanes-Lundback, B. Vaillancourt et al., 2017 Novel loci underlie natural variation in vitamin E levels in maize grain. Plant Cell 29(10): 2374–2392.

Diepenbrock, C. H., and M. A. Gore, 2015 Closing the divide between human nutrition and plant breeding. Crop Sci. 55: 1437–1448.

Elshire, R. J., J. C. Glaubitz, Q. Sun, J. A. Poland, K. Kawamoto et al., 2011 A robust, simple genotyping-by-sequencing (GBS) approach for high diversity species. PLoS ONE 6: e19379.

Estevez, J. M., A. Cantero, A. Reindl, S. Reichler and P. Leon, 2001 1-Deoxy-D-xylulose-5-phosphate synthase, a limiting enzyme for plastidic isoprenoid biosynthesis in plants. J. Biol. Chem. 276: 22901–22909.

Eroglu, A., and E. H. Harrison, 2013 Carotenoid metabolism in mammals, including man: formation, occurrence, and function of apocarotenoids. J. Lipid Res. 54(7): 1719–1730.

Ewels, P., M. Magnusson, S. Lundin, and M. Kaller, 2016 MultiQC: summarize analysis results for multiple tools and samples in a single report. Bioinformatics 32: 3047–3048.

Fang, H., X. Fu, Y. Wang, J. Xu, H. Feng et al., 2020 Genetic basis of kernel nutritional traits during maize domestication and improvement. Plant J. 101: 278–292.

Flint-Garcia, S. A., A. C. Thuillet, J. Yu, G. Pressoir, S. M. Romero et al., 2005 Maize association population: a high-resolution platform for quantitative trait locus dissection. Plant J. 44: 1054–1064.

Foudree, A., A. Putarjunan, S. Kambakam, T. Nolan, J. Fussell et al., 2012 The mechanism of variegation in *immutans* provides insight into chloroplast biogenesis. Front. Plant Sci. 3: 260.

Fu, J., Y. Cheng, J. Linghu, X. Yang, L. Kang et al., 2013 RNA sequencing reveals the complex regulatory network in the maize kernel. Nat. Commun. 4: 2832.

Fu, Z., Y. Chai, Y. Zhou, X. Yang, M. L. Warburton et al., 2013 Natural variation in the sequence of *PSY1* and frequency of favorable polymorphisms among tropical and temperate maize germplasm. Theor. Appl. Genet. 126: 923–935.

Gebremeskel, S., A. L. Garcia-Oliviera, A. Menkir, V. Adetimirin, and M. Gedil, 2018 Effectiveness of predictive markers for marker assisted selection of pro-vitamin A carotenoids in medium-late maturing maize (*Zea mays* L.) inbred lines. J. Cereal. Sci. 79: 27–34.

Gilmore, A. M., 1997 Mechanistic aspects of xanthophyll cycle-dependent photoprotection in higher plant chloroplasts and leaves. Physiol. Plantarum 99: 197–209.

Glaubitz, J. C., T. M. Casstevens, F. Lu, J. Harriman, R. J. Elshire et al., 2014 TASSEL-GBS: a high capacity genotyping by sequencing analysis pipeline. PLoS ONE 9: e90346.

Gonzalez-Jorge, S., P. Mehrshahi, M. Magallanes-Lundback, A. E. Lipka, R. Angelovici et al., 2016 *ZEAXANTHIN EPOXIDASE* activity potentiates carotenoid degradation in maturing seed. Plant Physiol. 171: 1837–1851.

Gonzalez-Jorge, S., S. H. Ha, M. Magallanes-Lundback, L. U. Gilliland, A. Zhou et al., 2013 Carotenoid cleavage dioxygenase4 is a negative regulator of beta-carotene content in Arabidopsis seeds. Plant Cell 25: 4812–4826.

Gore, M. A., J. M. Chia, R. J. Elshire, Q. Sun, E. S. Ersoz et al., 2009 A first-generation haplotype map of maize. Science 326: 1115–1117.

Graham, R. D., R. M. Welch and H. E. Bouis, 2001 Addressing Micronutrient Malnutrition Through Enhancing the Nutritional Quality of Staple Foods: Principles, Perspectives, and Knowledge Gaps. Adv. Agron. 70: 77–142.

Harjes, C. E., T. R. Rocheford, L. Bai, T. P. Brutnell, C. B. Kandianis et al., 2008 Natural genetic variation in *lycopene epsilon cyclase* tapped for maize biofortification. Science 319: 330–333.

Harker, M., and P. M. Bramley, 1999 Expression of prokaryotic 1-deoxy-D-xylulose-5-phosphatases in *Escherichia coli* increases carotenoid and ubiquinone biosynthesis. FEBS Lett. 448: 115–119.

Holland, J. B., W. E. Nyquist and C. T. Cervantes-Martínez, 2003 Estimating and interpreting heritability for plant breeding: an update in Plant Breeding Reviews, edited by J. Janick. John Wiley & Sons, Inc.

Hoopes, G. M., J. P. Hamilton, J. C. Wood, E. Esteban, A. Pasha, B. Vaillancourt, N. J. Provart, and C. R. Buell, 2018 An Updated Gene Atlas for Maize Reveals Organ-Specific and Stress-Induced Genes. Plant J. 97: 1154–1167.

Hung, H. Y., C. Browne, K. Guill, N. Coles, M. Eller et al., 2012 The relationship between parental genetic or phenotypic divergence and progeny variation in the maize nested association mapping population. Heredity 108: 490–499.

Jahns, P., and A. R. Holzwarth, 2012 The role of the xanthophyll cycle and of lutein in photoprotection of photosystem II. Biochim. Biophys. Acta 1817: 182–193.

Jia, K. P., L. Baz, and S. Al-Babili, 2018 From carotenoids to strigolactones. J. Exp. Bot. 69: 2189–2204.

Jin, X., C. Bai, L. Bassie, C. Nogareda, I. Romagosa et al., 2019 ZmPBF and ZmGAMYB transcription factors independently transactivate the promoter of the maize (*Zea mays*) β-carotene hydroxylase 2 gene. New Phytol. 222: 793–804.

Kandianis, C. B., R. Stevens, W. Liu, N. Palacios, K. Montgomery et al., 2013 Genetic architecture controlling variation in grain carotenoid composition and concentrations in two maize populations. Theor. Appl. Genet. 126: 2879–2895.

Kermode, A. R., 2005 Role of abscisic acid in seed dormancy. J. Plant Growth Regul. 24: 319–344.

Khoo, H. E., K. N. Prasad, K. W. Kong, Y. Jiang and A. Ismail, 2011 Carotenoids and their isomers: color pigments in fruits and vegetables. Molecules 16: 1710–1738.

Kim, D., G. Pertea, C. Trapnell, H. Pimentel, R. Kelley, and S. L. Salzberg, 2013 TopHat2: accurate alignment of transcriptomes in the presence of insertions, deletions and gene fusions. Genome Biol. 14: R36.

Kistler, L., S. Y. Maezumi, J. G. de Souza, N. A. S. Przelomşka, F. M. Costa et al., 2018 Multiproxy evidence highlights a complex evolutionary legacy of maize in South America. Science 362(6420): 1309–1313.

Koornneef, M., L. Bentsink and H. Hilhorst, 2002 Seed dormancy and germination. Curr. Opin. Plant Biol. 5: 33–36.

Krinsky, N. I., J. T. Landrum and R. A. Bone, 2003 Biologic mechanisms of the protective role of lutein and zeaxanthin in the eye. Annu. Rev. Nutr. 23: 171–201.

Krzywinski, M., J. Schein, I. Birol, J. Connors, R. Gascoyne et al., 2009 Circos: an information aesthetic for comparative genomics. Genome Res 19: 1639–1645.

Kurtz, S.: The Vmatch large scale sequence analysis software. 2019. URL http://www.vmatch.de.

Kutner, M. H., C. J. Nachtsheim, J. Neter and W. Li, 2004 Applied Linear Statistical Models. McGraw Hill Irwin, Boston.

Lee, M., N. Sharopova, W. D. Beavis, D. Grant, M. Katt et al., 2002 Expanding the genetic map of maize with the intermated B73 x Mo17 *(IBM) population*. Plant Mol. Biol. 48: 453–461.

Li, F., R. Vallabhaneni, J. Yu, T. Rocheford and E. T. Wurtzel, 2008 The maize phytoene synthase gene family: overlapping roles for carotenogenesis in endosperm, photomorphogenesis, and thermal stress tolerance. Plant Physiol. 147: 1334–1346.

Li, H., P. Bradbury, E. Ersoz, E. S. Buckler and J. Wang, 2011 Joint QTL linkage mapping for multiple-cross mating design sharing one common parent. PLoS ONE 6: e17573.

Li, L., H. Yuan, Y. Zeng and Q. Xu, 2016 Plastids and carotenoid accumulation. Subcell. Biochem. 79: 273–293.

Lu, S., Y. Zhang, K. Zhu, W. Yang, J. Ye et al., 2018 The citrus transcription factor CsMADS6 modulates carotenoid metabolism by directly regulating carotenogenic genes. Plant Physiol. 176: 2657–2676.

Martin, M., 2011 Cutadapt removes adapter sequences from high-throughput sequencing reads. EMBnet.journal 17: 10–12.

Mayer, M. P., P. Beyer and H. Kleinig, 1990 Quinone compounds are able to replace molecular oxygen as terminal electron acceptor in phytoene desaturation in chromoplasts of *Narcissus pseudonarcissus* L. Eur. J. Biochem. 191: 359–363.

McMullen, M. D., S. Kresovich, H. S. Villeda, P. Bradbury, H. Li et al., 2009 Genetic properties of the maize nested association mapping population. Science 325: 737–740.

Meenakshi, J. V., A. Banerji, V. Manyong, K. Tomlins, N. Mittal et al., 2012 Using a discrete choice experiment to elicit the demand for a nutritious food: willingness-to-pay for orange maize in rural Zambia. J. Health Econ. 31: 62–71.

Menkir, A., B. Maziya-Dixon, W. Mengesha, T. Rocheford, and E. Oladeji, 2017 Accruing genetic gain in pro-vitamin A enrichment from harnessing diverse maize germplasm. Euphytica 213: 105.

Neter, J., M. H. Kutner, C. J. Nachtsheim and W. Wasserman, 1996 Applied Linear Statistical Methods. Irwin, Chicago.

Norris, S. R., T. R. Barrette and D. DellaPenna, 1995 Genetic dissection of carotenoid synthesis in Arabidopsis defines plastoquinone as an essential component of phytoene desaturation. Plant Cell 7: 2139–2149.

Ogut, F., Y. Bian, P. J. Bradbury and J. B. Holland, 2015 Joint-multiple family linkage analysis predicts within-family variation better than single-family analysis of the maize nested association mapping population. Heredity 114: 552–563.

Owens, B. F., A. E. Lipka, M. Magallanes-Lundback, T. Tiede, C. H. Diepenbrock et al., 2014 A foundation for provitamin A biofortification of maize: genome-wide association and genomic prediction models of carotenoid levels. Genetics 198: 1699–1716.

Pixley K, N. P. Palacios, R. Babu, R. Mutale, and E. Simpungwe, 2013 Biofortification of maize with provitamin A carotenoids. *In* Carotenoids, Human Health and Nutrition, S. A. Tanumihardo, ed. (Springer Science + Business Media, New York), pp. 271–292.

Portwood, II, J. P., M. R. Woodhouse, E. K. Cannon, J. M. Gardiner, L. C. Harper et al., 2019 MaizeGDB 2018: the maize multi-genome genetics and genomics database. Nucleic Acids Res. 47(D1): D1146–D1154.

Prasanna, B. M., N. Palacios-Rojas, F. Hossain, V. Muthusamy, A. Menkir et al., 2020 Molecular breeding for nutritionally enriched maize: status and prospects. Front. Genet. 10: 1392.

Quackenbush, F. W., J. G. Firch, W. J. Rabourn, M. McQuistan, E. N. Petzold et al., 1961 Analysis of carotenoids in corn grain. J. Agr. Food Chem. 9: 132–135.

Quinlan, R. F., M. Shumskaya, L. M. Bradbury, J. Beltran, C. Ma et al., 2012 Synergistic interactions between carotene ring hydroxylases drive lutein formation in plant carotenoid biosynthesis. Plant Physiol. 160: 204–214.

R Core Team (2018). R: A language and environment for statistical computing. R Foundation for Statistical Computing, Vienna, Austria. URL https://www.R-project.org/.

Ramstein, G. R., S. J. Larsson, J. P. Cook, J. W. Edwards, E. S. Ersoz et al., 2020 Dominance effects and functional enrichments improve prediction of agronomic traits in hybrid maize. Genetics 215(1): 215–230.

Rice, A. L., K. P. West Jr., and R. E. Black, 2004. Vitamin A deficiency. *In* Comparative Quantification of Health Risks: Global and Regional Burden of Disease Attributable to Selected Major Risk Factors, M. Ezzati, A. D. Lopez, A. Rodgers, and C. J. L. Murray, eds. Volume 1. (World Health Organization, Geneva, Switzerland), pp. 211–256.

Rodermel, S., 2002 Arabidopsis variegation mutants. Arabidopsis Book 1: e0079.

Rodriguez-Villalon, A., E. Gas and M. Rodriguez-Concepcion, 2009 Phytoene synthase activity controls the biosynthesis of carotenoids and the supply of their metabolic precursors in dark-grown Arabidopsis seedlings. Plant J. 60: 424–435.

Rodgers-Melnick, E., P. J. Bradbury, R. J. Elshire, J. C. Glaubitz, C. B. Acharya et al., 2015 Recombination in diverse maize is stable, predictable, and associated with genetic load. P. Natl. Acad. Sci. USA 112(12): 3823–3828.

Rosso, D., A. G. Ivanov, A. Fu, J. Geisler-Lee, L. Hendrickson et al., 2006 IMMUTANS does not act as a stress-induced safety valve in the protection of the photosynthetic apparatus of Arabidopsis during steady-state photosynthesis. Plant Physiol. 142: 574–585.

Rosso, D., R. Bode, W. Li, M. Krol, D. Saccon et al., 2009 Photosynthetic redox imbalance governs leaf sectoring in the Arabidopsis thaliana variegation mutants *immutans*, *spotty*, *var1*, and *var2*. Plant Cell 21: 3473–3492.

Saltzman, A., E. Birol, H. E. Bouis, E. Boy, F. F. De Moura et al., 2013 Biofortification: progress toward a more nourishing future. Glob. Food Sec. 2: 9–17.

SAS Institute Inc. 2011. Base SAS® 9.3 Procedures Guide: Statistical Procedures. Cary, NC: SAS Institute Inc.

Schwarz, Gideon, 1978 Estimating the dimension of a model. Ann. Statist. 6(2): 461–464.

Scrucca, L., M. Fop, T. B. Murphy, and A. E. Raftery, 2016 mclust 5: clustering, classification and density estimation using Gaussian finite mixture models. The R Journal 8(1): 289–317.

Shahbazi, M., M. Gilbert, A. M. Laboure and M. Kuntz, 2007 Dual role of the plastid terminal oxidase in tomato. Plant Physiol. 145: 691–702.

Stahl, W., and H. Sies, 2005 Bioactivity and protective effects of natural carotenoids. Biochim. Biophys. Acta 1740: 101–107.

Stanley, L. E., B. Ding, W. Sun, F. Mou, C. Hill et al., 2020 A tetratricopeptide repeat protein regulates carotenoid biosynthesis and chromoplast development in monkeyflowers (*Mimulus*). Plant Cell 32: 1536–1555.

Suwarno, W. B., K. V. Pixley, N. Palacios-Rojas, S. M. Kaeppler and R. Babu, 2014 Formation of heterotic groups and understanding genetic effects in a provitamin A biofortified maize breeding program. Crop Sci. 54: 14–24.

Suwarno, W. B., K. V. Pixley, N. Palacios-Rojas, S. M. Kaeppler and R. Babu, 2015 Genome-wide association analysis reveals new targets for carotenoid biofortification in maize. Theor. Appl. Genet. 128: 851–864.

Swarts, K., R. M. Gutaker, B. Benz, M. Blake, R. Bukowski et al., 2017 Genomic estimation of complex traits reveals ancient maize adaptation to temperate North America. Science 357: 512–515.

Taleon, V., L. Mugode, L. Cabrera-Soto, and N. Palacios-Rojas, 2017 Carotenoid retention in biofortified maize using different post-harvest storage and packaging methods. Food Chem. 232: 60–66.

Tan, B.-C., J.-C. Guan, S. Ding, S. Wu, J. W. Saunders et al., 2017 Structure and origin of the *White Cap* locus and its role in evolution of grain color in maize. Genetics 206: 135–150.

Tan, B.-C., L. M. Joseph, W.-T. Deng, L. Liu, Q.-B. Li et al., 2003 Molecular characterization of the Arabidopsis 9-cis epoxycarotenoid dioxygenase gene family. Plant J. 35: 44–56.

Tian, F., P. J. Bradbury, P. J. Brown, H. Hung, Q. Sun et al., 2011 Genome-wide association study of leaf architecture in the maize nested association mapping population. Nat. Genet. 43: 159–162.

Tian, L., V. Musetti, J. Kim, M. Magallanes-Lundback and D. DellaPenna, 2004 The *Arabidopsis LUT1* locus encodes a member of the cytochrome P450 family that is required for carotenoid E-ring hydroxylation activity. Proc. Natl. Acad. Sci. 101: 402–407.

Trapnell, C., A. Roberts, L. Goff, G. Pertea, D. Kim, D. R. Kelley, H. Pimentel, S. L. Salzberg, J. L. Rinn, and L. Pachter, 2012 Differential gene and transcript expression analysis of RNA-seq experiments with TopHat and Cufflinks. Nat. Protoc. 7: 562–578.

Tuteja, N., 2007 Abscisic acid and abiotic stress signaling. Plant Signal Behav. 2: 135–138.

Valdar, W., C. C. Holmes, R. Mott, and J. Flint, 2009 Mapping in structured populations by resample model averaging. Genetics 182: 1263–1277.

Vallabhaneni, R., L. M. Bradbury and E. T. Wurtzel, 2010 The carotenoid dioxygenase gene family in maize, sorghum, and rice. Arch. Biochem. Biophys. 504: 104–111.

Vogel, J. T., B.-C. Tan, D. R. McCarty and H. J. Klee, 2008 The carotenoid cleavage dioxygenase 1 enzyme has broad substrate specificity, cleaving multiple carotenoids at two different bond positions. J. Biol. Chem. 283: 11364–11373.

Wallace, J. G., P. J. Bradbury, N. Zhang, Y. Gibon, M. Stitt et al., 2014 Association mapping across numerous traits reveals patterns of functional variation in maize. PLoS Genet. 10: e1004845.

Weber, E. J., 1987 Carotenoids and tocols of corn grain determined by HPLC. J. Am. Oil Chem. Soc. 64: 1129–1134.

Weir, B. S., 1996 Genetic data analysis II. Sinauer Associates.

Welch, R. M., and R. D. Graham, 2004 Breeding for micronutrients in staple food crops from a human nutrition perspective. J. Exp. Bot. 55: 353–364.

West, K.P., and I. Darnton-Hill. 2008. Vitamin A deficiency. In: R.D. Semba and M. Bloem, editors, Nutrition and health in developing countries, 2nd ed. The Humana Press, Inc., Totowa, NJ. p. 377–433.

West, K. P., Jr., 2002 Extent of vitamin A deficiency among preschool children and women of reproductive age. J. Nutr. 132: 28575–28665.

Wetzel, C. M., and S. R. Rodermel, 1998 Regulation of phytoene desaturase expression is independent of leaf pigment content in *Arabidopsis thaliana*. Plant Mol. Biol. 37: 1045–1053.

Wetzel, C. M., C.-Z. Jiang, L. J. Meehan, D. F. Voytas and S. R. Rodermel, 1994 Nuclear-organelle interactions: the *immutans* variegation mutant of *Arabidopsis* is plastid autonomous and impaired in carotenoid biosynthesis. Plant J. 6: 161–175.

Wong, W. L., X. Su, X. Li, C. M. G. Cheung, R. Klein et al., 2014 Global prevalence of age-related macular degeneration and disease burden projection for 2020 and 2040: a systematic review and meta-analysis. The Lancet Global Health 2: e106–e116.

Wu, D., D. A. Wright, C. Wetzel, D. F. Voytas and S. Rodermel, 1999 The *IMMUTANS* variegation locus of Arabidopsis defines a mitochondrial alternative oxidase homolog that functions during early chloroplast biogenesis. Plant Cell 11: 43–55.

Yan, J., C. B. Kandianis, C. E. Harjes, L. Bai, E.-H. Kim et al., 2010 Rare genetic variation at *Zea mays crtRB1* increases beta-carotene in maize grain. Nat. Genet. 42: 322–327.

Yu, J., J. B. Holland, M. D. McMullen and E. S. Buckler, 2008 Genetic Design and Statistical Power of Nested Association Mapping in Maize. Genetics 178: 539–551.

Zhang, L., X. Zhang, X. Wang, J. Xu, M. Wang et al., 2019 SEED CAROTENOID DEFICIENT functions in isoprenoid biosynthesis via the plastid MEP pathway. Plant Physiol. 179: 1723–1738.

Zhu, C., S. Naqvi, J. Breitenbach, G. Sandmann, P. Christou et al., 2008 Combinatorial genetic transformation generates a library of metabolic phenotypes for the carotenoid pathway in maize. Proc. Natl. Acad. Sci. 105: 18232–18237.

Zunjare, R. U., R. Chhabra, F. Hossain, A. Baveja, V. Muthusamy et al., 2018 Molecular characterization of 5’ UTR of the *lycopene epsilon cyclase* (*lcyE*) gene among exotic and indigenous inbreds for its utilization in maize biofortification. 3 Biotech. 8: 75.

